# Multiomics Characterization of Potential Therapeutic Vulnerabilities in Low-grade Serous Ovarian Carcinoma

**DOI:** 10.1101/2020.06.18.135061

**Authors:** Raunak Shrestha, Marta Llaurado Fernandez, Amy Dawson, Joshua Hoenisch, Stanislav Volik, Yen-Yi Lin, Shawn Anderson, Hannah Kim, Anne Haegert, Shane Colborne, Brian McConeghy, Robert H. Bell, Sonal Brahmbhatt, Gabriel E. DiMattia, Stephane Le Bihan, Gregg B. Morin, Colin C. Collins, Mark S. Carey

## Abstract

**Background:** Low-grade serous ovarian carcinoma (LGSOC) is a rare tumor subtype with high case fatality rates. As such, there is a pressing need to develop more effective treatments using newly available preclinical models for therapeutic discovery and drug evaluation. Here, we use a multiomics approach to interrogate a collection of LGSOC patient-derived cell lines to elucidate novel biomarkers and therapeutic vulnerabilities.

**Methods:** Fourteen LGSOC cell lines were interrogated using whole exome sequencing, RNA sequencing, and mass spectrometry-based proteomics. Somatic mutation, copy-number aberrations, gene and protein expression were analyzed and integrated using different computational approaches. LGSOC cell line data was compared to publicly available LGSOC tumor data (AACR GENIE cohort), and also used for predictive biomarker identification of MEK inhibitor (MEKi) efficacy. Protein interaction databases were evaluated to identify novel therapeutic targets.

**Results:** *KRAS* mutations were exclusively found in MEKi-sensitive and *NRAS* mutations mostly in MEKi-resistant cell lines. Analysis of COSMIC mutational signatures revealed distinct patterns of nucleotide substitution mutations in MEKi-sensitive and MEKi-resistant cell lines. Deletions of *CDKN2A/B* and *MTAP* genes (chromosome 9p21) were much more frequent in cell lines than tumor samples and possibly represent key driver events in the absence of KRAS/NRAS/BRAF mutations. For *in-vitro* MEKi efficacy prediction, proteomic data provided better discrimination than gene expression data. Condensin, MCM, and RFC protein complexes were identified as potential treatment targets in MEKi-resistant cell lines.

**Conclusions:** Our LGSOC cell lines are representative models of the most common molecular aberrations found in LGSOC tumors. This study highlights the importance of using proteomic data in multiomics assessment of drug prediction and identification of potential therapeutic targets. CDKN2A/B and MTAP deficiency offer an opportunity to find synthetically lethal candidates for novel treatments. Multiomics approaches are crucial to improving our understanding of the molecular aberrations in LGSOC, establishing effective drug prediction programs and identifying novel therapeutic targets in LGSOC.

## Background

Low-grade serous ovarian carcinoma (LGSOC) is a rare subtype of epithelial ovarian cancer with a poor prognosis. Women with LGSOC are usually diagnosed with advanced-stage disease and only 10-20% are alive 10 years after diagnosis [1,2]. Research on LGSOC is challenging due to its low prevalence, uncertain etiology, and the limited availability of research models. However, in the last 5-10 years investigators have elucidated many key genomic aberrations, leading to major advancements in the molecular characterization and classification of LGSOC [3,4]. In contrast to other ovarian cancer subtypes, LGSOC are genetically characterized by high frequency of oncogenic mutations in *KRAS, NRAS*, and *BRAF* (20-40%, 7-26%, 5-33%, respectively) [5–7], a very low prevalence of *TP53* mutations (<8%) [8,9], frequent copy-number deletion of *CDKN2A* resulting in loss of tumor suppressor protein p16 (15-53%) [10,11], and high expression of estrogen receptor (ER) and progesterone receptor (PR) (>90% and >50%, respectively) [12–14]. More recently, less frequent mutations in *USP9X* (15%) and *EIF1AX* (8%) have been described in a small proportion of LGSOC tumors [10,15].

Traditionally, the therapeutic management of different ovarian cancer histotypes has been similar as it is now appreciated that different ovarian histological subtypes represent different diseases [16]. Consequently, the most common treatment for LGSOC is based on cytoreductive surgery followed by platinum/paclitaxel chemotherapy. However, unlike the most common subtype of epithelial ovarian cancer (high-grade serous ovarian carcinoma or HGSOC), LGSOC is relatively resistant to standard chemotherapy [2,17]. Anti-estrogen therapy is also a common therapy though response rates for the treatment of recurrent disease patients are low (9-14%) [18]. In this context, the identification of oncogenic mutations affecting MAPK genes (RAS/RAF/MEK) led to the evaluation of targeted agents such as MEK inhibitors (MEKi) in clinical trials of patients with LGSOC. Recently, a study using the MEKi trametinib for the treatment of relapsed LGSOC, demonstrated a statistically significant improvement in progression free survival (PFS) when compared to standard of care therapies. Trametinib monotherapy resulted in an improvement in PFS of almost 6 months (13 vs 7.2 months) with objective response rates of 26% (compared to 6% in the standard treatment arm). While promising, these results demonstrated that benefit of trametinib therapy is restricted to a subgroup of patients. Thus, additional effective therapies are needed.

Advancements in the treatment of rare cancers such as LGSOC rests on a better understanding of their unique molecular features and how they can be exploited for therapeutic gain. Unfortunately, results from genomic precision oncology clinical trials have shown that only 5-7% of cancer patients derive benefit [19,20]. However, in this small fraction of patients, response rates were over 50% and durable response durations of 30 months were observed. To accelerate progress in precision oncology, future research must go beyond the genomic aberrations found in tumors and explore the more complex aspects of tumor biology. Thus, multiomics tumor profiling with data integration analysis [21] provides a new window of opportunity for the identification of molecular disease drivers leading to more successful drug therapies. This approach can be extremely useful in the study of rare cancers such as LGSOC, where there is an urgent need for more effective therapies.

In the last few years, we have established a large collection of cancer cell line models from advanced/recurrent LGSOC patients [22]. These models have since then been used for pre-clinical drug testing and biomarker discovery in several studies [22–24]. With the introduction of MEKi therapy for LGSOC, there is a pressing need to identify biomarkers that predict tumor response. These predictive biomarkers will not only allow us to identify those patients that are more likely to benefit from MEKi therapy, and through further research identify novel treatment targets for the resistant patient population [24].

In this study, we performed an integrative analysis of the genomes, transcriptomes, and proteomes of 11 clinically characterized LGSOC cell line models with the goal to identify novel tumor vulnerabilities for the development of future drug treatments. Here, using multiomics approach, we aim to: (1) characterize the molecular complexity of our LGSOC models, (2) elucidate novel potential drivers associated with LGSOC disease, and (3) identify molecular biomarkers that predict response to MEKi therapy.

## Methods

### Tumor samples and clinical information

Advanced or recurrent LGSOC samples (tumor and ascites) were obtained from the OvCaRe gynecologic tumor bank (Vancouver General Hospital (VGH), British Columbia Cancer Agency, and the John and Mary Knight Translational Ovarian Cancer Research Unit (London Regional Cancer Program, London, Ontario, Canada). Tumor bank protocols, cell line derivation, and all research relating to this study was approved by institutional human ethics review boards at BCCA (H14-02859), the University of British Columbia (UBC; R05-0119), and the University of Western Ontario (HSREB 12668E). Clinical information was extracted retrospectively from patient records. Tumor histology was confirmed by a gynecological pathologist.

### Patient-derived LGSOC cell lines

Low-grade serous ovarian cancer patient-derived cell lines were established through continuous in-vitro culture of patient material obtained through OvCaRe or the John and Mary Knight Translational Ovarian Cancer Research Unit (cell line iOvCa241) tumor banks [22,24]. For details, see Supplementary Methods.

### Whole exome sequencing

DNA was extracted from LGSOC cell lines using an AllPrep DNA/RNA Mini Kit (Qiagen, Toronto, ON), and from matched patient buffy coat samples using a DNeasy Blood and Tissue Kit (Qiagen, Toronto, ON), according to the manufacturer’s protocol. DNA concentration was quantified using a Qubit^*®*^ dsDNA HS Assay (Thermo Scientific, Burlington, ON, Canada). About 0.5 μg of genomic DNA was fragmented by hydrodynamic shearing (Covaris, Inc., Woburn, MA, USA) to generate 150-280 bp fragments. After end repair, fragments were ligated to Illumina barcoded adapters, cleanup, amplified by polymerase chain reaction (PCR) to enrich for adapter ligated fragments, and controlled for quality. The library was then enriched by liquid phase hybridization using Agilent SureSelect XT Human All Exon v6 (Agilent Technologies, CA, USA) following the manufacturer’s recommendations. The captured library was amplified by PCR using indexing primers, cleanup and quality assessment was done with the TapeStation 4200 (Agilent). Libraries are submitted to PE100 sequencing on Illumina HiSeq4000.

### Somatic variant calling

Sequence alignment and mutation calling were performed in Partek Flow environment (© Partek Inc). Sequence reads were aligned to the GRCh38/hg38 human genome build using bwa 0.7.2 [25]. Variants were called using Strelka 1.0.15 [26] for all cell lines except for CL-02 (lacking buffy coat sample). CL-02 variant calling was performed using LoFreq 2.1.3.a [27]. The called variants were annotated using the Annovar software [28]. Annotated calls were then filtered to show only protein-changing single nucleotide variations (SNVs) that were present in cell line DNA at allele frequencies (AF) greater than 0.1 and coverage higher than 16x. For CL-02, all calls not present in dbSNP (version 138) were retained, while of the calls that were present in dbSNP, calls with (average heterozygosity + aveHet standarderror) less than 0.1 were retained. These were additionally filtered using the same criteria as for the Strelka calls.

### Copy number aberration (CNA) calls

Data analysis was performed using Nexus Copy Number Discovery Edition Version 9.0 (BioDiscovery, Inc., El Segundo, CA). Samples were processed using the Nexus NGS functionality with the FASST2 segmentation. The log ratio thresholds for single copy gain and single copy loss were set at +0.18 and - 0.18, respectively. The log ratio thresholds for gain of 2 or more copies and for a homozygous loss were set at +0.6 and -1.0, respectively. Tumor sample Binary alignment map (BAM) files were processed with corresponding normal tissue BAM files. Probes were normalized to the median.

### Whole transcriptome sequencing (RNA-seq)

Total RNA from was isolated in accordance with the mirVana miRNA Isolation kit (ThermoFisher). Strand specific RNA sequencing was performed on quality controlled high RIN value (>7) RNA samples (Bioanalyzer Agilent Technologies). In brief, 200ng of total RNA was first treated to remove the rRNA and then purified using the Agencourt RNA Clean XP Kit (Beckman Coulter) prior to analysis with the Agilent RNA 6000 Pico Chip to confirm rRNA removal. Next, the rRNA-depleted RNA was fragmented and converted to cDNA. Subsequent steps include end repair, addition of an ‘A’ overhang at the 3’ end, ligation of the indexing-specific adaptor, followed by purification with Agencourt Ampure XP beads. The strand specific RNA library prepared using TruSeq stranded mRNA kit (Illumina). Libraries were quality checked and sized with a TapeStation 4200 (Agilent Technologies), then quantified using the KAPA Library Quantification Kit for Illumina platforms (Kapa Biosystems). Libraries were submitted to PE75 sequencing on Illumina NextSeq500. About 20-50 million sequencing reads per library was analyzed for each sample.

### Transcriptome (RNA-seq) quantification

RNA-seq sequence alignment and transcript/gene quantification were performed in Partek Flow environment (© Partek Inc). The RNA-seq reads were first trimmed to remove low quality bases from the 3’ end. The trimmed reads were then mapped to the GRCh38/hg38 human reference genome and transcriptome (RefSeq release 83) using splice-aware aligner STAR (2.5.3a) [29]. Based on the reads that can only be mapped to a single genomic location, the transcript/gene expression quantification was performed using HTSeq-count [30]. Cross-sample normalization of expression values were done by DESeq [31].

### Proteomics analysis using mass spectrometry

Seven LGSOC cell lines were selected for global proteome analysis (**Supplementary Table 1**) in triplicates using 3 separately cultured samples for each. Cultures of 5 different HGSOC cell lines (OVCAR8, OVSAYO, CAOV3, OVCAR4, and VOA2900) were also used as a control. VOA2900 is a HGSOC cell line developed by the ovarian cancer research program at VGH. Cells were collected by trypsinization and washed with 1X dPBS. A minimum of 1.0×10^6^ cells per cell line were used in these analyses. Cells pellets were lysed in 200 uL of a buffer containing guanidine hydrochloride (4M), HEPES (pH 8.5, 50 mM), 2-chloroacetamide (40 mM), tris(2-carboxyethyl) phosphine (10 mM), ethylenediaminetetraacetic acid (5 mM), 1x phosphatase inhibitor, and 1x protease inhibitor. Bead beating of the cells was performed in Lysing Matrix D tubes on a FastPrep24 (6 M/s for 45 s, 60 s rest, 6 M/s for 45 s). Cell lysate was then heated at 95 C for 15 min with 1200 rpm mixing. 50 ug of each sample was diluted 10x in 50 mM HEPES pH 8.5 (concentration determined using by BCA assay) and digested using a 50:1 ratio of protein:trypsin/Lys-C mixture at 37 C overnight with shaking at 1200 rpm. A small aliquot of each LGSOC sample was mixed and analyzed as LGSOC-mixed control. The five HGSOC samples were mixed and used as a HGSOC-mixed control sample. An aliquot of all the LGSOC and HGSOC samples were mixed and used as an all sample pooled internal standard. Tandem mass tag (TMT10) labeling was performed by adding 100 ug TMT label in acetonitrile and incubating at room temperature for thirty minutes twice. Excess label was quenched with 10 uL of 1 M glycine. Individual TMT channels were combined and the volume of the combined TMT labeled sample was reduced to 10-20% of the original volume. Three 10-plex batches were prepared, where each batch contained one replicate of each of the LGSOC samples, the LGSOC-mixed control, the HGSOC mixed control, and the all sample pooled internal standard. The combined samples were fractionated by high pH reverse-phase high performance liquid chromatography (HPLC) in a gradient of acetonitrile and 20 mM ammonium bicarbonate in water into 48 individual fractions. These 48 fractions were concatenated into 12 fractions by combining every twelfth fraction, each of which was analyzed by low pH nanoLC mass spectrometry (MS) using a MS/MS/MS (MS^3^) method on a Thermo Scientific Easy nLC coupled to a ThermoFisher Orbitrap Fusion MS. The Uniprot Human Proteome was used as a reference proteome [32]. Only peptides that were identified in all three batches were retained. ComBat, a function from *sva* R-package [33], was used to adjust for batch effects using an empirical Bayes adjustment. Detection of differentially expressed proteins from the collected peptide data was determined by the Probe-level Expression Change Averaging (PECA) in R [34].

### Mutational signature analysis

We used deconstructSigs [35], a multiple regression approach to statistically quantify the contribution of mutational signature for each tumor. The 30 mutational signatures were obtained from the COSMIC mutational signature database [36]. In brief, deconstructSigs attempts to recreate the mutational pattern using the trinucleotide mutation context from the input sample that closely resembles each of the 30 mutational signatures from COSMIC mutational signature database (v2 - March 2015). In this process, each mutational signature is assigned a weight normalize between 0 to 1 indicating its contribution. Only those mutational signatures with a weight more than 0.06 were considered for analysis. We further combined (or grouped) these 30 mutational signatures based on the similarity of their mutational features or functional associations. For details, see Supplementary Methods.

### Prioritization of driver genes using HIT’nDRIVE

HIT’nDRIVE [37] measures the potential impact of genomic aberrations on changes in the global expression of other genes/proteins which are in close proximity in a gene/protein-interaction network. It then prioritizes those aberrations with the highest impact as cancer driver genes. Both non-silent somatic mutation calls and CNA gain or loss were independently collapsed in gene-patient alteration matrix with binary labels. mRNA and protein expression values were used to derive expression-outlier gene-patient outlier matrix using GESD test. STRING ver10 [38] protein-interaction network was used to compute pairwise influence value between the nodes in the interaction network. We integrated these genome and transcriptome as well as genome and proteome data using HIT’nDRIVE. First we integrated genome and transcriptome data. Here we ran HIT’nDRIVE separately using SNV-mRNA expression data and CNA-mRNA expression data. For this we used the parameters: **α**=0.9, **β**=0.7, and **γ**=0.7. Next, we integrated genome and proteome data. Here we ran HIT’nDRIVE separately using SNV-protein expression data and CNA-protein expression data. For this we used the parameters: **α**=0.9, **β**=0.7, and **γ**=0.8. We used IBM-CPLEX as the Integer Linear Programming solver. The results were later combined to downstream analysis.

### Differential expression analysis

Differential expression analysis of MEKi-response phenotypes was performed by applying linear empirical Bayes model using “sva” R-package [33]. Gene (mRNA) and protein expression data of the MEKi-response phenotypes were used for this purpose. We used the following threshold values to select genes and proteins for MEKi response analysis and downstream pathway analysis: for mRNA expression data, *pvalue* ≤ 0.05 and |*foldchange*| *>* 1.03, and for protein expression data, *pvalue* ≤ 0.05 and |*foldchange*| *>* 1.5. Furthermore, to identify differentially expressed protein complex, we used Wilcoxon rank-sum test on the average protein expression profiles of protein complex members. Protein complexes with *pvalue* ≤ 0.05 and |*foldchange*| *>* 1 were selected differential protein complex analysis.

### Pathway enrichment analysis

The differentially expressed set of genes or proteins were tested for enrichment against gene sets of KEGG pathways present in Molecular Signature Database (MSigDB) v6.0 [39]. A hypergeometric test based over-representation analysis was used for this purpose (https://github.com/raunakms/GSEAFisher). A cut-off threshold of false discovery rate (FDR) ≤ 0.01 was used to obtain the significantly enriched pathways. Only pathways that are enriched with at least four differentially expressed genes were considered for further analysis. To calculate the pathway activity score, the expression dataset was transformed into standard normal distribution using “inverse normal transformation” method. This step is necessary for a fair comparison between the expression-values of different genes. For each sample, the pathway activity score is the mean expression level of the differentially expressed genes linked to the enriched pathway. The KEGG signaling pathways and associated expression profile were visualized using *pathview* R-package [40].

### External datasets

We utilized DNA sequencing datasets of publicly available patient LGSOC cohort from the American Association for Cancer Research (AACR) project Genomics Evidence Neoplasia Information Exchange (GENIE) [41]. The dataset consisted of a total of 122 LGSOC tumors (both primary and metastatic tumors). We used somatic mutation and copy number aberration profiles from the dataset. AACR project GENIE Data: Version 5.0 was downloaded from (SynapseID: syn7222066) https://www.synapse.org/#!Synapse:syn7222066.

### Protein complex co-expression score

For a given protein complex, its co-expression score is computed as the average Pearson correlation coefficient of all pairwise protein-protein interactions (as represented in STRING ver 10 database [38]) in the complex. The co-expression scores were separately calculated using mRNA expression data (*R*_*mRNA*_) and protein expression data (*R*_*protien*_). Protein complexes with (|*R*_*mrna*_ − *R*_*protien*_| ≤ 0.05) are defined as correlated protein complexes. For this analysis, we considered the comprehensive resource of mammalian protein complexes (CORUM) protein complexes [42] consisting of at least 4 proteins.

## Results

### Landscape of somatic mutations in LGSOC

To investigate the landscape of somatic mutation changes in LGSOC cell lines, we performed high-coverage whole exome sequencing (WES) of 14 LGSOC cell lines and 10 matched normal samples from 11 independent patients (3 patients had 2 consecutive tumors cultured and included in this study) (**Supplementary Table 1**). We refer to this cohort of LGSOC cell lines as VGH cohort. We achieved a mean coverage of 106x for normal samples and 107x for tumor samples (**Supplementary Table 2**). Altogether, we detected 1893 highly confident non-silent mutations in the protein-coding regions in 14 cell lines (**Supplementary Table 3-4**).

As described in our recent study [24], non-synonymous mutations in *KRAS* were found in all four MEKi-sensitive LGSOC cell lines (CL-01, CL-02, CL-14, and CL-15) (**Fig. 1A**). Two of them (CL-01 and CL-14) harbored *KRAS*^*G12D*^ mutations while the other two (CL-02 and CL-15) harbored *KRAS*^*G12V*^ mutations (**Fig. 1B-C**). The variant allele frequency (VAF) of these *KRAS* mutations were over 70% in CL-14, CL-15, and CL-01 and that in CL-02 was 55% (**Supplementary Figure 1**) indicating that these mutations are highly likely to be clonal in origin. Interestingly, all mutations detected in *KRAS* were the glycine residue at position 12. *NRAS* was mutated in one MEKi-sensitive cell line and three MEKi-resistant cell lines. *NRAS*^*Q61R*^ mutation was specific to the three MEKi-resistant cell lines (CL-03, CL-04, CL-10) whereas, *NRAS*^*C118Y*^ mutation was present in one MEKi-sensitive cell line (CL-14) with a co-occurring *KRAS* mutation. The VAF of *NRAS* mutations ranged between 48-62% also indicating that these mutations are highly likely to be clonal. (**Supplementary Figure 1**). A *BRAF*^*D594G*^ mutation was present in only one cell line (CL-09) (**Fig. 1B**). We further evaluated somatic mutations in a large cohort of LGSOC tumors (n=97, includes both primary and metastatic tumors, but the tumor stages information are not available) in AACR project GENIE [41]. All mutations identified in *KRAS, NRAS*, and *BRAF* in the VGH cohort were localized in the same amino-acid positions as that reported in the GENIE cohort (**Fig. 1C**). However, *BRAF*^*D594G*^ mutation identified in the VGH cohort was not present in tumors GENIE cohort but one tumor sample harbored a mutation in the same amino acid position (*BRAF*^*D594N*^) (**Supplementary Figure 2**). Ten cell lines, including all MEKi-sensitive cases, had at least one mutated gene in the MAPK pathway (**Supplementary Figure 3A**). In the GENIE cohort, *RAS/RAF m*utations were prevalent (63%) and were mutually exclusive (**Supplementary Figure 2**).

**Figure 1.**
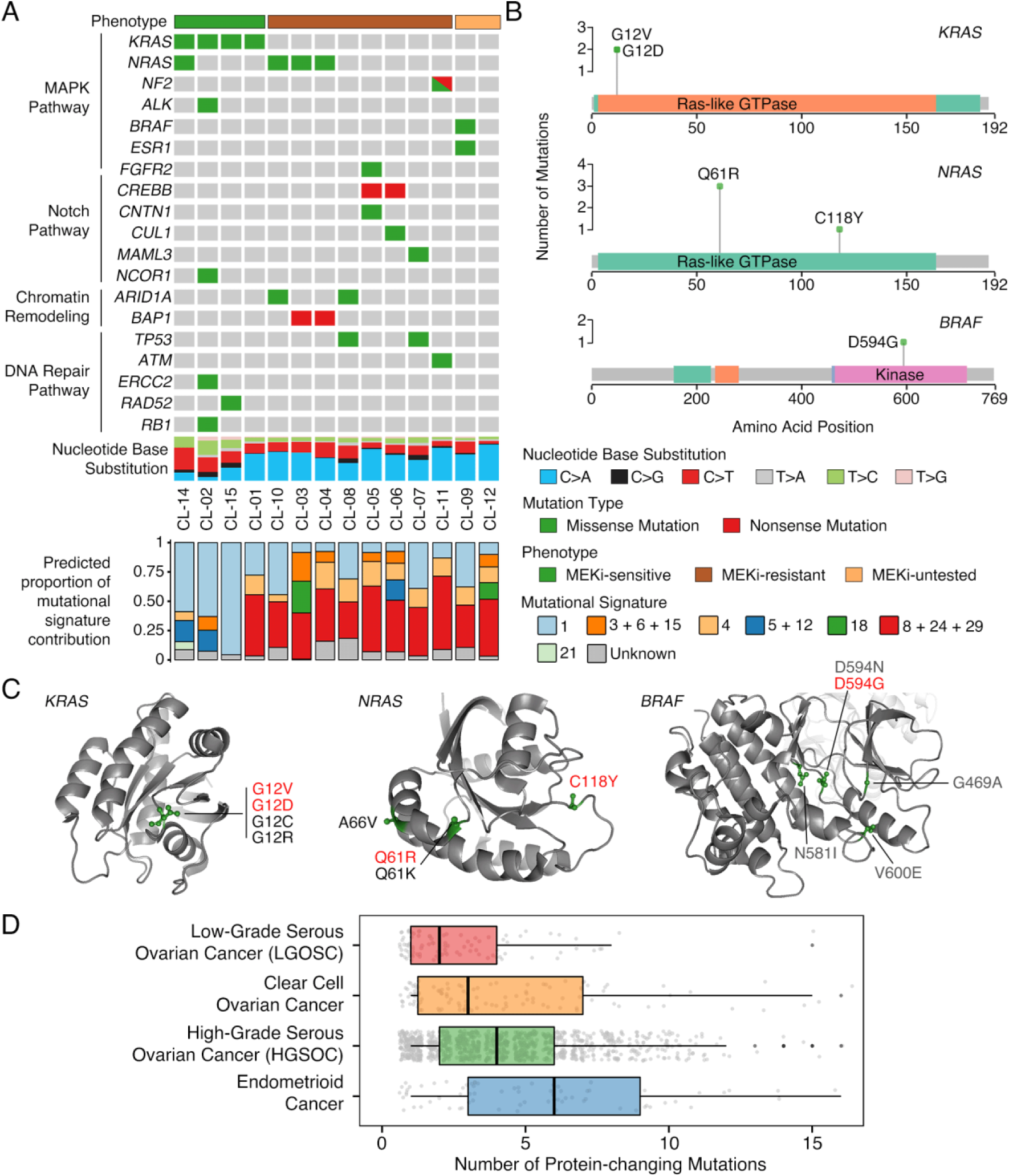
Landscape of somatic mutation in LGSOC. (A) Status of somatic mutations in fourteen LGSOC cell lines grouped by genes in major cancer pathways - MAPK pathway, Notch pathway, chromatin remodeling, and DNA repair pathway. The bottom panel represents the proportional contribution of different COSMIC mutational signatures per sample. (B) Plots showing mutation distribution and the protein domains for the corresponding mutated protein. (C) 3D protein structure of KRAS, NRAS, and BRAF with mutations identified in LGSOC cases in both VGH cohort (LGSOC cell lines, highlighted in red) and GENIE cohorts (LGSOC tumors) were highlighted. (D) Box plot showing the comparison of the tumor mutation burden in LGSOC with that of major subtypes of ovarian cancer. We leveraged the tumors included in the GENIE cohort for this analysis. Each dot in the plot represents a tumor sample of respective ovarian cancer subtype.

Apart from mutations in MAPK signaling pathway axis, other sparsely mutated genes identified in the LGSOC cell lines were involved in Notch pathway, chromatin remodeling, and DNA repair (**Fig. 1A, Supplementary Figure 3A**). Notably, *TP53*^R234H^ mutations were observed in two MEKi-resistant cell lines (CL-07 and CL-08) both with VAF above 96% indicating their clonal origin (**Fig 1A, Supplementary Figure 1**). While mutations in *TP53* are rare in LGSOC tumors (8% frequency) [9,43], these cell lines (which were derived from two sequential tumor samples corresponding to the same patient) had been confirmed to be LGSOC by pathology review [44]. Furthermore, 3% of tumors in the GENIE cohort had *TP53* mutations. Notch pathway genes were mutated in 5 cell lines; mostly in MEKi-resistant cases. Signaling pathways such as PI3K, chromatin remodeling, MYC, TP53, and NRF2 were mostly mutated in MEKi-resistant cell lines (**Supplementary Figure 3A**). Similarly, different subsets of LGSOC tumors in GENIE cohort also harbored mutations in DNA repair pathway, chromatin remodeling pathway, Notch pathway, and TP53 pathway (**Supplementary Figure 3B**).

Using the software deconstructSigs [35], we evaluated the characteristic mutation patterns in our patient-derived cell lines against the mutational signatures obtained from the COSMIC mutational signature database [36]. Analyses of the mutation patterns in the cell lines revealed strong enrichment of C>T transition nucleotide substitution mutations in MEKi-sensitive cell lines whereas MEKi-resistant and MEKi-untested cell lines were enriched in both C>T (transition) as well as C>A (transversion) nucleotide substitution mutations (**Fig. 1A and Supplementary Figure 4**). Moreover, GENIE cohort of LGSOC tumors also demonstrated strong enrichment of C>T and C>A nucleotide substitution mutations. We found mutational signature 1, predominantly related to age-related mutagenesis, to be operative in MEKi-sensitive cell lines (**Fig. 1A**). On the other hand, mutational signatures 4, 8, 24, and 29 that are associated with transcriptional strand bias for C>A mutations were operative in MEKi-resistant and MEKi-untested cell lines. Interestingly, mutational signatures 3, 6, and 15 that are associated with DNA mismatch repair were also operative in five MEKi-resistant and MEKi-untested cell lines. Clinical annotation of these samples revealed that MEKi-sensitive cell lines were associated with primary tumors (rather than recurrent tumors) that had received a lower number of therapy regimens prior collection than the MEKi-resistant associated cultures. Interestingly, MEKi-sensitive cell lines, despite being associated with older patients (average 65.5 vs. 51.7 years), their estimated overall survival rates were found to be longer (average 6.5 vs. 4.4 years) (**Supplementary Table 1**). The presence of serous borderline or micropapillary tumor patterns, as well as the time to recurrence were not found to be associated with the two distinct MEKi response phenotype observed in our LGSOC cell lines.

Finally, using the data available from the GENIE cohort, we compared the exonic tumor mutation burden (TMB) in the four major subtypes of ovarian cancer (HGSOC, LGSOC, clear cell ovarian cancer, and endometrioid ovarian cancer) in the GENIE cohort. Compared to the other subtypes, LGSOC tumors have a low TMB (**Fig. 1D**).

### Landscape of copy number aberrations in LGSOC

Copy number aberrations (CNAs) were determined using the whole exome sequence reads. We identified 6480 CNAs across 14 LGSOC cell lines analyzed (**Supplementary Table 5**). Copy number changes for all cell lines are shown in **Fig. 2A**, and those for individual cell lines in **Fig. 2B.** As previously described, loss of chromosome 9p and gains in chromosomes 8, 12 and 20 are frequent [22]. Overall, homozygous copy loss of *MTAP* was observed in 13 out of 14 LGSOC cell lines and its loss of heterozygosity (LOH) was observed in 1 cell line (**Fig. 2C**). Similarly, homozygous copy loss of *CDKN2A/B* was observed in 12 out of 14 LGSOC cell lines and its LOH was observed in 2 cell lines. These copy number changes were associated with loss of p16 expression by western blot (data available upon request).

**Figure 2.**
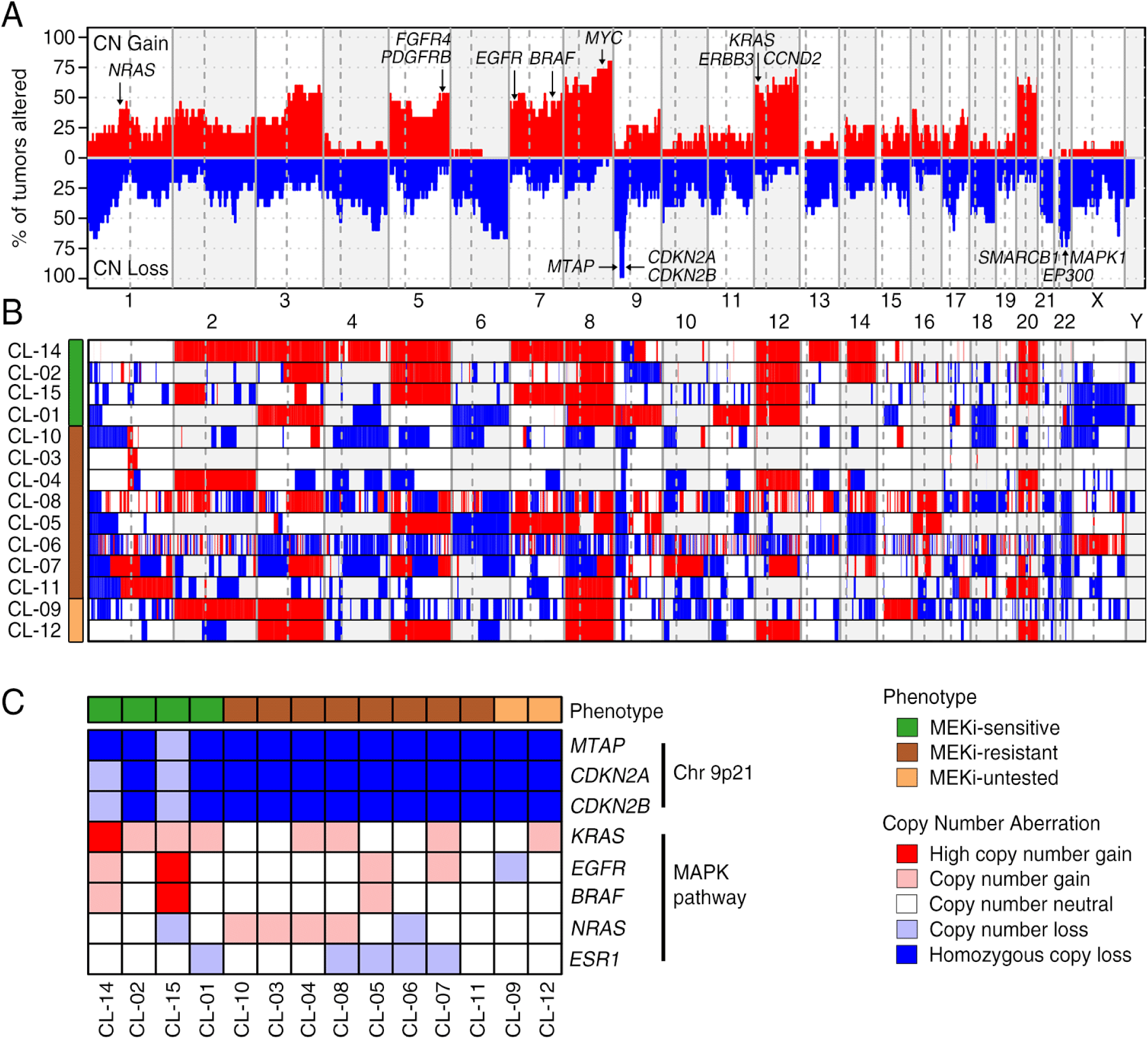
Landscape of copy number aberrations in LGSOC. (A) Aggregate copy number alteration profile of LGSOC cell lines analyzed shown by each chromosome region. Red indicates copy number gain and blue indicates copy number loss. Important genes with copy number changes are highlighted. (B) Copy number alteration profile per sample analyzed. (C) Copy number status of some selected genes in chromosome 9p21 and MAPK pathway.

Overall GISTIC analysis of focal amplifications and deletion revealed several recurrent events containing known oncogenic drivers (**Supplementary Table 6**). These include amplifications of *PDGFRB* (5q32), *FGFR4* (5q35), *MYC* (8q24), *CCND2* (12p13) along with deletions of *CDKN2A* (9p21), *MAPK1* (22q11), and *SMARCB1* (22q11). All four MEKi-sensitive LGSOC cell lines, which were *KRAS* mutant, also harbored copy-number gains affecting *KRAS*. CNA were identified in different genes targeting key oncogenic pathways such as MAPK, NOTCH, PI3K, Hippo, Wnt, TGF-beta signaling pathways, chromatin remodeling, DNA repair pathways, and cell cycle process (**Supplementary Figure 5**). We observed copy number deletions of few genes in Chromatin Remodeling, NOTCH, and Wnt signaling pathways enriched in MEKi-resistant cell lines. Interestingly, MEKi-sensitive cell lines are for the most part copy neutral on chromosome 1, 9p, and 10 and also tend to have chromosome 5, 8, and 12 copy number gains.

### Multiomics approaches to identify LGSOC drivers and predict MEKi-response

#### Identification of driver genes in LGSOC using HIT’nDRIVE

Using our recently developed computational algorithm HIT’nDRIVE [37], we identified driver genes in these LGSOC cell lines. We first ran HIT’nDRIVE using SNV-mRNA expression data and CNA-mRNA expression data. HIT’nDRIVE analysis prioritized 17 unique driver genes in 8 LGSOC cell lines for which matched genome and transcriptome data were available (**Fig. 3A** and **Supplementary Table 7**). Similarly, we also ran HIT’nDRIVE using SNV-protein expression data and CNA-protein expression data. This analysis identified 19 unique driver genes in 7 LGSOC cell lines for which matched genome and proteome data were available (**Fig. 3B** and **Supplementary Table 7**).

**Figure 3.**
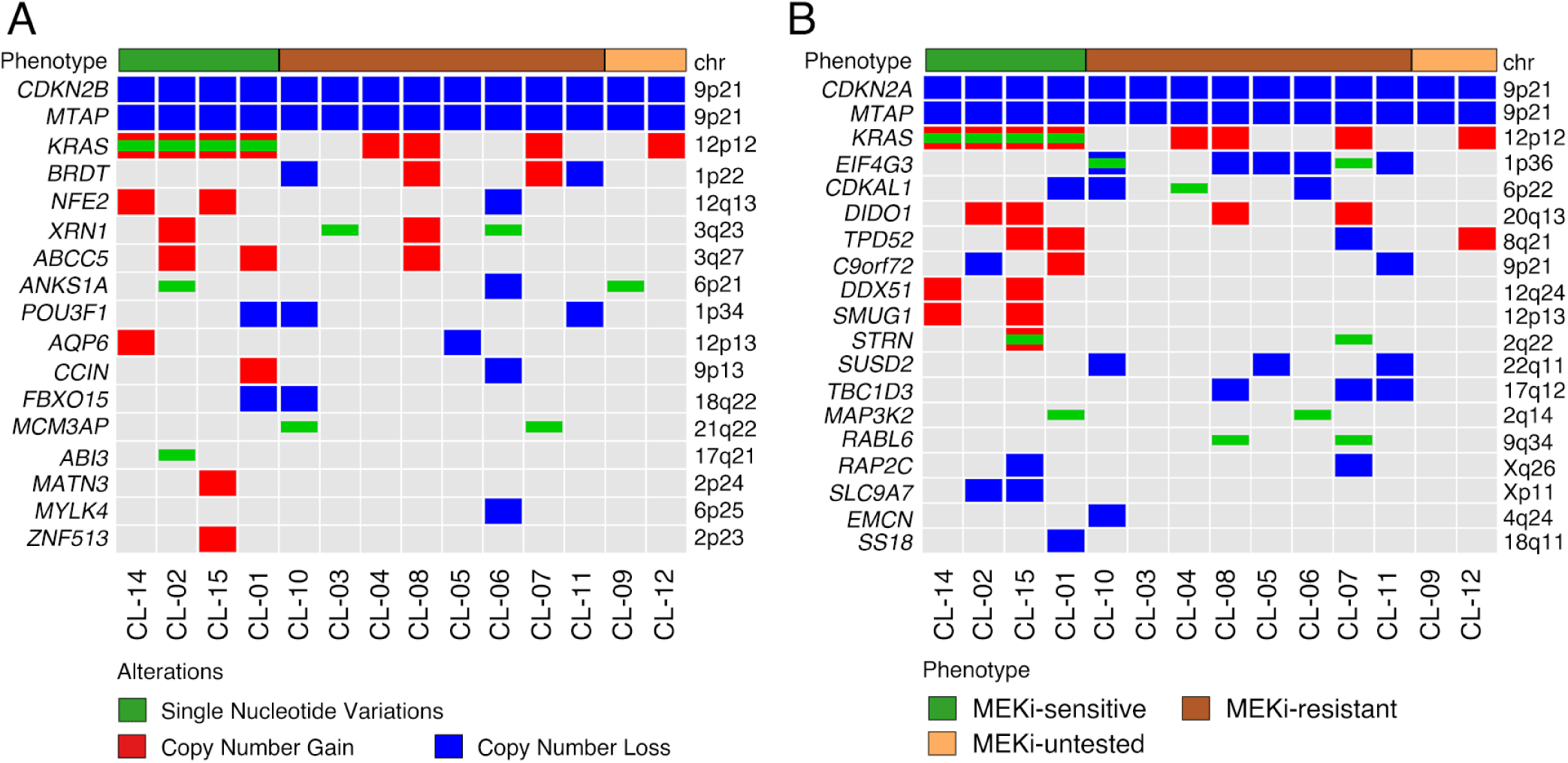
Driver genes of LGSOC identified by HIT’nDRIVE. (A-B) These oncoplots represent the genomic alterations in the driver genes of LGSOC identified by HIT’nDRIVE. To identify the driver genes, SNV and CNA data were used along with (A) mRNA expression data, and (B) protein expression data. Mutation and copy number aberration data were analyzed separately using HIT’nDRIVE. We note that: HIT’nDRIVE analysis were performed in (A) 8 LGSOC cell lines with matched genome and mRNA expression data and (B) 7 LGSOC cell lines with matched genome and protein expression data available. i.e. we could not use 6 cell lines without mRNA-expression profiles and 7 cell lines without protein-expression profile for HIT’nDRIVE analysis. However, in these cell lines not used for HIT’nDRIVE analysis, if the predicted driver genes resulting from other cell lines were present, we represented them as a part of the above figure.

Both HIT’nDRIVE analyses using mRNA and protein expression data identified genes in chromosome 9p21, *MTAP, CDKN2A*, and *CDKN2B*, as the most common driver genes among all LGSOC cell lines (**Fig. 3A-B**). For both CDKN2A and MTAP, the mRNA and protein expression profiles (available for 7 cell lines) were found to correlate and to be expressed at very low levels. Analysis of CNA in the LGSOC tumors in the GENIE cohort revealed that loss of *CDKN2A/B* and *MTAP* was present in about 31.5% (17 out of 54) of all tumors, with few homozygous deletions 7.4% (4 out of 54) (**Supplementary Table 8**).

Using HIT’nDRIVE CNA-mRNA expression analysis copy number gains that were shared by more than one cell line include *ABCC5, XRN1, BRDT*, and *NFE2*. Similarly, HIT’nDRIVE CNA-protein expression analysis revealed CNA gain of *DIDO1, TPD52, DDX51*, and *SMUG1* in more than one cell line. Interestingly, *NEF2, DDX51*, and *SMUG1* are located on chromosome 12q indicating potential role of this region in LGSOC progression. Furthermore, HIT’nDRIVE CNA-protein expression analysis prioritized mutation and loss of gene EIF4G3 specific to MEKi-resistant phenotype.

#### Evaluating gene and protein expression to predict MEKi-response

Next, we performed total RNA-seq (Illumina NextSeq500) on 8 LGSOC cell lines with known sensitivity to MEKi treatment [22]. We detected 14.3 million reads on an average and over 27700 unique expressed genes across all samples. Principal components analysis (PCA) using the RNA-seq profiles obtained from each sample revealed that, except for the sample CL-01, all other MEKi-sensitive samples were clustered together whereas MEKi-resistance samples had high variance and thus were spread along the principal component axes (**Fig. 4A**). This suggests that the MEKi-sensitive samples are transcriptionally very similar to each other, whereas the MEKi-resistant samples are very diverse.

**Figure 4.**
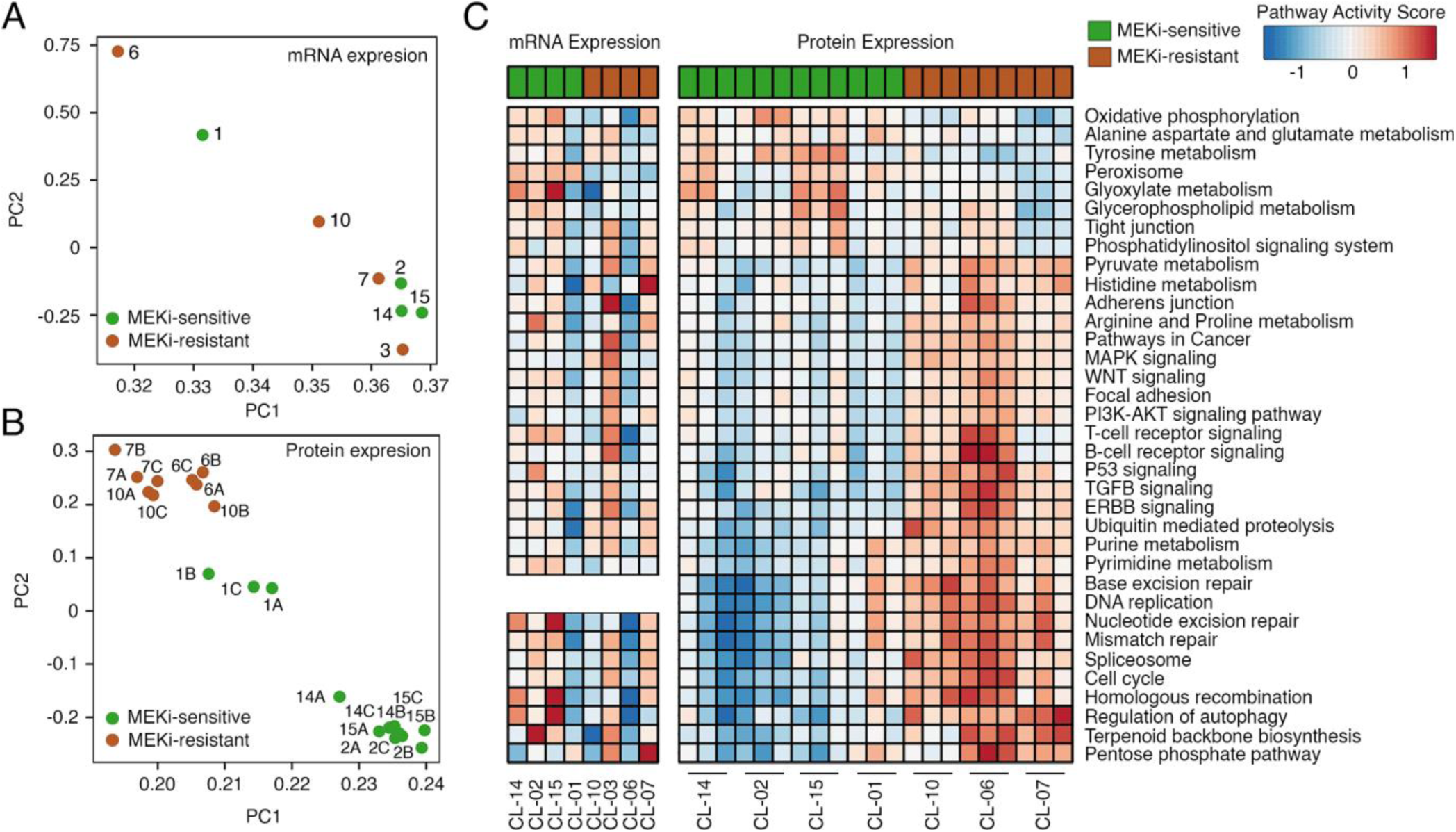
Characterization of transcriptome and proteome profile of MEKi response phenotypes. (A-B) Principal component analysis (PCA) of LGSOC using (A) 4710 most variable genes in mRNA expression profiles and (B) 1278 most variable proteins in proteome expression profiles. See **Methods** section for details. (C) Pathways enrichment of differentially expressed genes/proteins between LGSOC MEKi treatment phenotypes obtained using mRNA expression and protein expression (biological triplicates).

Next, we performed proteome analysis by mass spectrometry on 7 LGSOC cell lines in triplicate and identified 5110 unique proteins. The PCA analysis on the mass spectrometry protein expression data also revealed two distinct clusters for MEKi-sensitive and MEKi-resistant samples (**Fig. 4B**). This demonstrates that the protein expression profiles of LGSOC cell lines discriminates the MEKi-sensitive and MEKi-resistant phenotype better than their mRNA expression profiles. Similar to the PCA in mRNA expression data, the CL-01 sample in PCA in the protein expression data was quite distinct as compared to the rest of the MEKi-sensitive samples.

We then identified differentially expressed transcripts and proteins between MEKi-sensitive and MEKi-resistant phenotype (see **Methods** section). A total of 1,231 and 1,202 differentially expressed genes (in mRNA expression profiles) and proteins (in protein expression profiles) were respectively identified, out of which 113 gene/protein pairs were detected in both datasets (**Supplementary Figure 6A and Supplementary Table 9-10**). Similar to the results obtained from the PCA analysis, the expression profiles of the differentially expressed proteins discriminates the MEKi-sensitive and MEKi-resistant phenotype better than that of the differentially expressed genes (mRNA) (**Supplementary Figure 6B-C**). The differentially expressed genes/proteins between MEKi-sensitive and MEKi-resistant LGSOC cell lines included *EGFR, PRKCA, MAPK4, NF2, MTOR, MTAP, ATM, TGFB2*, and *BRCA2*.

To identify signaling pathways dysregulated by the differentially expressed genes/proteins between MEKi-sensitive and MEKi-resistant cell lines, we performed gene set enrichment analysis (see **Methods** section for details) (**Supplementary Table 11-12**). Although, distinct sets of genes (in the mRNA expression profile) and proteins (in the protein expression profile) were found to be differentially expressed between the MEKi-response phenotypes, intriguingly, we observed similar sets of differentially expressed signaling pathways enriched between the MEKi-response phenotypes between the two datasets (**Fig. 4C-D**). We found that majority of the proteins in the signaling pathways such as MAPK pathway, Wnt pathway, P53 pathway, DNA Replication, DNA Repair pathway, Cell cycle process, and Amino-acid metabolism were found to be highly expressed in MEKi-resistant as compared to MEKi-sensitive cell lines. On the other hand, we found majority of the proteins in Oxidative Phosphorylation and Tight Junction pathways to be highly expressed in the MEKi-sensitive lines. Similar to the observation made in **Fig. 4A-B** (and **Supplementary Fig. 7**), the pathway level analysis also indicates that the protein expression profiles separate the MEKi response phenotypes much more distinctly than that from mRNA expression data. Thus, in the downstream analysis we mostly present analysis based on protein expression profiles.

#### Protein complexes characterizing MEKi response phenotypes

As a novel discovery approach, we sought to identify large protein complexes characterizing the differences in the MEKi response phenotypes, we leveraged a curated set of core protein complexes from the CORUM database [42]. To identify protein complexes that characterize the MEKi response phenotypes, we performed Wilcoxon rank-sum test on the average protein expression profiles of CORUM protein complex members (see **Methods** section). In MEKi-resistant cell lines, we identified protein complexes such as BRCC complex, p130-ER-alpha complex, VEGFR2 complex, Condensin complex, minichromosome maintenance (MCM) complex and replication factor C (RFC) complex to have high protein-expression of their complex members in MEKi-resistant cell lines (**Fig. 5A-B and Supplemental Table 13**). In MEKi-sensitive cell lines, we found F1F0-ATP synthase complex and RNA-induced silencing complex (RISC) to have high protein-expression of their complex members.

**Figure 5.**
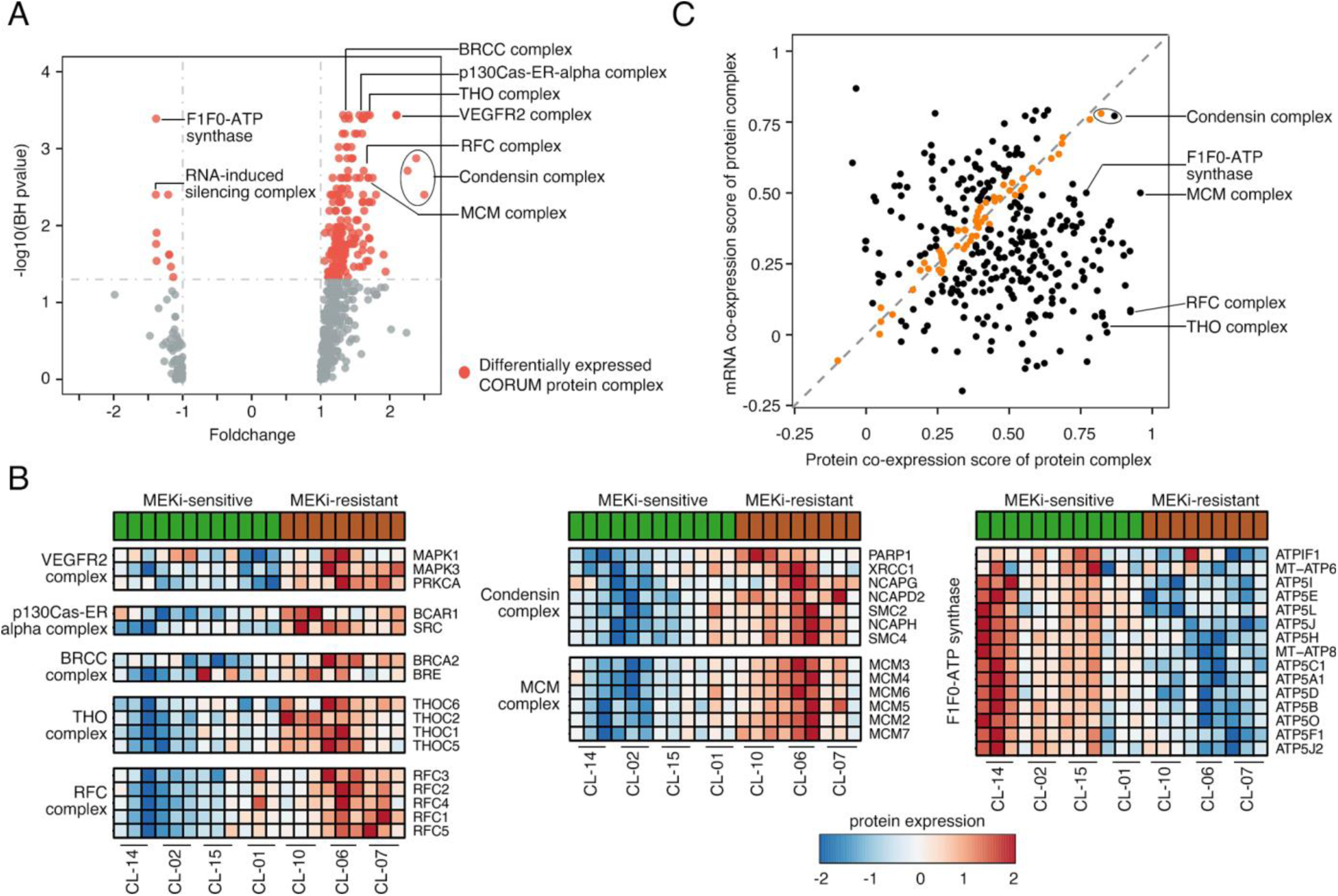
Protein complexes characterizing MEKi response phenotypes. (A) Volcano plot showing the differentially expressed CORUM protein complexes between MEKi sensitive and resistant LGSOC cell lines. Each dot represents a CORUM protein complex. Differentially expressed protein complexes are represented as red colored dots. The expression fold-change is calculated relative to MEKi-resistant cell lines. (B) Heatmap of protein expression profiles of CORUM protein complexes that are differentially expressed between the MEKi response phenotypes across LGSOC cell lines. (C) Scatter plot showing the correlation between mRNA co-expression score and protein co-expression score of CORUM protein complexes. Each dot in the figure represents a protein complex and the correlated protein complexes are highlighted in orange color. p130Cas-ER-alpha complex: p130Cas-ER-alpha-cSrc-kinase-PI3-kinase p85-subunit complex; VEGFR2 complex: VEGFR2-S1PR1-ERK1/2-PKC-alpha complex.

Furthermore, to identify protein complexes whose both transcript and protein of its members were co-expressed, we measured their co-expression score (see **Methods** section). We found 56 (or of 343) protein complexes with correlated mRNA and protein co-expression (**Figure. 5C and Supplementary Table 14**). We also found 28 protein complexes with high mRNA co-expression score (*R*_*mRNA*_ > 0.5) as well as high protein co-expression score (*R*_*protein*_ > 0.5). This included the Condensin complex which was found to have strong co-expression scores. MCM and RFC complexes on the other had strong protein co-expression but weak mRNA co-expression.

#### Dysregulation of MAPK signaling pathway that may influence MEKi response

With the knowledge that MEKi treatment shows activity in some patients with incurable LGSOC disease, we further investigated MAPK pathway activation in more detail. First, we correlated mRNA and protein expression fold change of genes/proteins involved in this pathway (**Fig. 6A**). In MEKi-resistant cell lines, genes such as PRKCA, HSPA2, TGFB2, FGF3, TP53, and FLNC were highly upregulated in MEKi-resistant samples in both mRNA and protein expression data, and that these findings were highly correlated (**Fig. 6B**). On the other hand, MAPK13, HSPB1, and CACNA2D1 were upregulated in MEKi-sensitive cell lines. Furthermore, we found that the RAS-RAF-MEK-ERK axis was highly upregulated in the MEKi-resistant cell lines as compared to the MEKi-sensitive cell lines. (**Supplementary Figure 8**).

**Figure 6.**
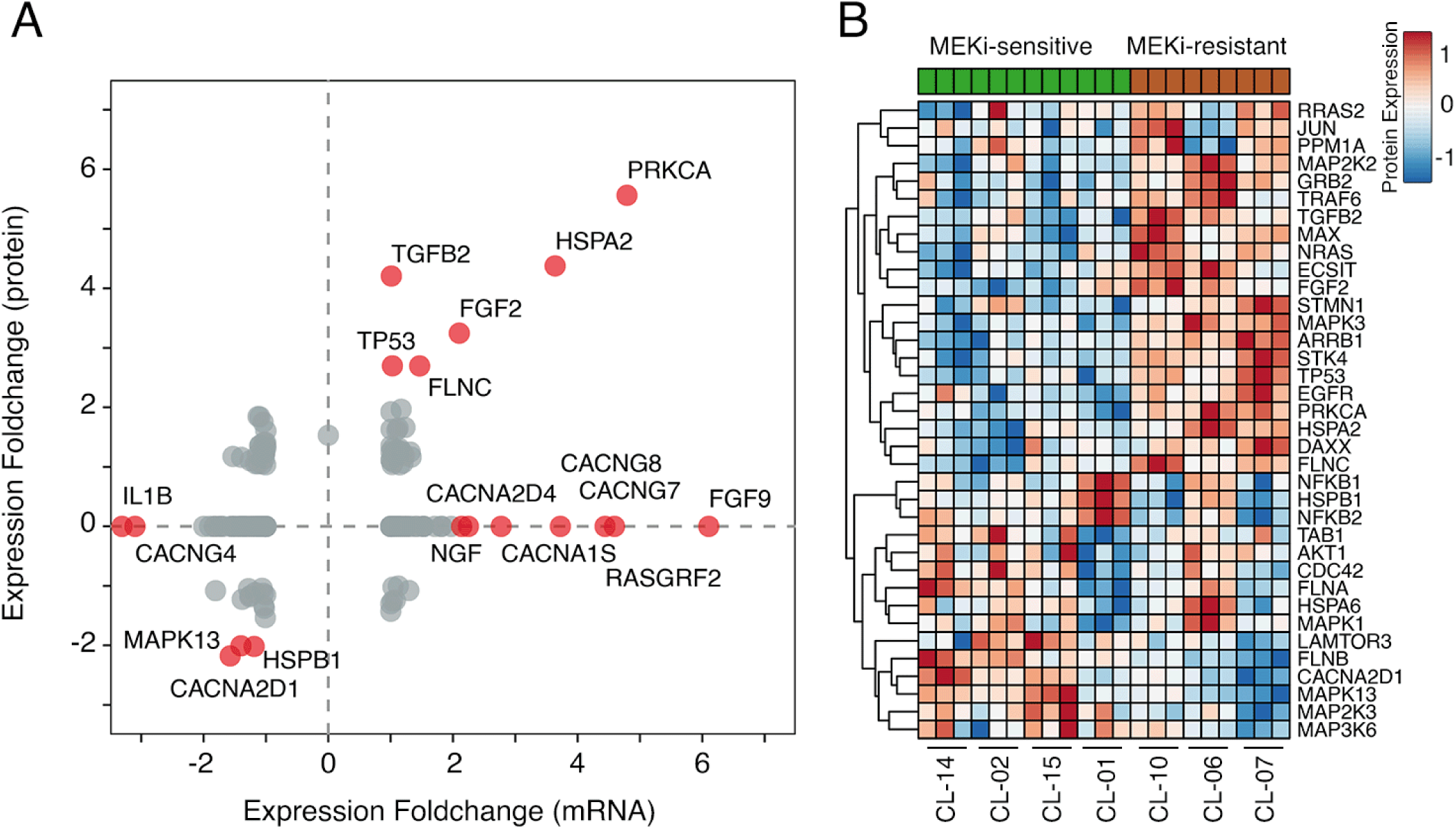
Proteome expression profile of MAPK signaling pathway characterizes response to MEKi treatment in LGSOC. (A) mRNA/Protein expression fold-change of genes or proteins involved in MAPK signaling pathway. Each dot represents a gene/protein. The expression fold-change is calculated relative to MEKi-resistant cell lines. The dots highlighted in red color are those genes/proteins with higher than 2 fold-change expression between the MEKi-resistant vs MEKi-sensitive LGSOC samples. (B) Heatmap of protein expression profiles of those proteins involved in MAPK signaling pathway.

## Discussion

Fatality rates in women with LGSOC are high due to a lack of effective treatments. Our study is the first to use a multiomics approach to characterize of the genome, transcriptome [45–47], and proteome [48–52] profiles of 14 patient-derived cell lines. These cell line models closely recapitulate the genomic abnormalities depicted in LGSOC tumors from the GENIE cohort [24] and other previous studies [5–7], and highlight common somatic mutations known to involve the MAPK (RAS/RAF/MEK) pathway [10,41,53]. LGSOC cell lines were characterized by a strong enrichment of C>T and C>A nucleotide substitution mutations, a predominance of *KRAS* mutations, and CNA’s that are characteristic of those described in previous reports of LGSOC [10,11]. Both LGSOC cell lines and tumors were characterized by a lower tumor mutation burden than other ovarian cancer subtypes. Our findings also support earlier observations that *BRAF* mutations are less frequent than *RAS* mutations in advanced/recurrent LGSOC cases [10,54,55].

Mutational signatures associated with LGSOC have not been previously described. Though these signatures have provided a remarkable basis for characterizing patterns of genomic disruption resulting from etiological factors in cancer generally [56], they may also be of importance as predictive biomarkers. However, more work needs to be done to identify if mutational signature might also have important implications for tumor classification and treatment management. Both LGSOC tumors and cell lines were found to be associated with transcriptional bias for C>A transversion nucleotide substitution mutations in LGSOC, indicating guanine damage that is being repaired by transcription-coupled nucleotide excision repair. Interestingly mutational signatures also distinguished MEKi response phenotypes in LGSOC cell lines. In particular, mutational signatures 1, 5, and 12 were associated with MEKi-sensitive cell lines, while signatures 8, 4, and 29 were much more prevalent in MEKi-resistant cell lines. Mutational signature 1, which is associated with age, correlated with older patients.

Loss of the chromosome 9p21 tumor suppressor locus (including *CDKN2A* and *MTAP*) is the most common deletion event across many human cancer types [57–61]. Both LGSOC cell lines and tumors had deletions affecting these genes. However, we noted an important difference in the frequency of these events. As previously described, LGSOC cell lines (86%; 12/14) had focal/homozygous deletions in chromosome 9p21, confirmed by a low CDKN2A/p16 expression. In contrast, LGSOC tumors from the GENIE project harbored heterozygous deletions of 9p21 in only 31.5-53% of cases. Thus, there appears to be a selective advantage for establishing primary cultures/cell lines of LGSOC with 9p loss. Interestingly, absence of p16 expression had been previously associated with an unfavorable prognosis in LGSOC [10,62] suggesting a potential value for the use of cell cycle inhibitors (CDK4) in LGSOC.

MTAP is a key enzyme in the methionine salvage pathway and is frequently co-deleted with *CDKN2A* due to its chromosomal proximity [63,64]. From large scale short hairpin RNA-mediated screens across multiple human cancer cell lines, it is known that viability of *MTAP*-deficient cancer cells is impaired by depletion of the protein arginine methyltransferase PRMT5 [58,63]. Inhibitors of PRMT5 may be synthetically lethal strategy for this dysregulated metabolic state and are of interest in *MTAP* and *CDKN2A/B*-deleted tumors. The potential therapeutic value of MTAP and CDKN2A deficiency in LGSOC remains to be explored.

Our study describes several different techniques to evaluate multiomics data for the identification of disease drivers and MEKi-response prediction. Recently, Schutte *et al*. showed integration of genomic and transcriptomic data significantly outperformed less comprehensive analysis in the identification of clinically relevant molecular alterations [65]. The combination of DNA and RNA-based analyses have shown to serve as mutual controls for verifying potential findings [21]. As LGSOC is a rare cancer, model systems and patient samples are limited. Thus, efficient analytical tools and large datasets are needed to reduce false discovery rates and improve prediction accuracy. HIT’nDRIVE analysis exploits the power of analyzing multiomics data and can also be used to enhance discovery in rare cancers [66]. Using HIT’nDRIVE analysis we identified novel potential driver genes (*KRAS, MTAP*, and *CDKN2A/B*) in LGSOC cell lines. In recent clinical trial with LGSOC patients, *KRAS* mutation were found to be associated with better response to MEK inhibition with significant improvement in PFS with binimetinib when compared to physician choice of chemotherapy [67]. Other single case reports have shown similar findings [68,69]. While promising results of MEKi therapy for the treatment of LGSOC patients have been demonstrated in recent studies [70,71], biomarkers that predict treatment efficacy are largely unknown.

Here using LGSOC cell lines, we demonstrated that proteomic profiles were better at discriminating distinct MEKi-response phenotypes than transcriptomic profiles, and found that protein alterations affecting TP53, cell cycle and DNA-repair pathways are common in MEKi-resistant cell lines. These pathways should be further studied for their potential therapeutic value. Proteomic technologies are useful for drug prediction [72]. The addition of proteomic-based analysis facilitates the understanding of the biological implications of genomic change and can be used as discovery tools for both drug prediction and identification of novel therapeutic targets. We found considerable consistency in the identification of differential signaling pathway activity between MEKi-response phenotypes using proteomic and transcriptomic data, strongly suggesting that these pathways play a role in MEKi resistance. Recent developments in analytical approaches using machine learning also show promise in improving multiomics drug prediction [73].

MEKi will soon be commonly prescribed, for the treatment of LGSOC, as an alternative to the standard chemotherapy and anti-hormone therapy in patients with recurrent disease [74]. To identify robust biomarkers that predict MEKi response, we used transcript/protein expression to evaluate MAPK pathway regulation. MAPK signaling feedback loops play a key role in MEKi resistance [75]. Both PI3K-AKT-mTOR pathway (**Supplementary Figure 9**) and MAPK pathway activity are upregulated in MEKi-resistant cell lines. The most differentially expressed protein identified in the MAPK pathway analysis was protein kinase C alpha (PRKCA or PKC-ɑ). We previously identified and validated both PKC-ɑ and EGFR as biomarkers of MEKi resistance using reverse phase protein array (RPPA) [24]. This observation supports the utility of multiomics analysis approaches for predictive biomarker discovery as we used a completely different proteomic technologies and dataset for discovery. This and other novel predictive biomarkers of MEKi efficacy have been described and some of these biomarkers could be considered for validation in future clinical trials.

While MEKi-sensitive LGSOC lines depend on MAPK pathway activation for survival (as shown by the lethal effects of MEKi treatment in-vitro), the importance of MAPK in MEKi-resistant lines remains unclear. In the absence of protein phosphorylation data, we cannot assess the extent of MAPK signaling activation in the MEKi-resistant cell lines. However, a more comprehensive evaluation of the function (activator/inhibitor) and activation status (phosphorylated/not phosphorylated) of the protein candidates identified in here will allow researchers to clarify if MEKi-resistant lines will benefit of drug combinations targeting different proteins within the MAPK pathway (e.g. MEK plus BRAF inhibitors), or if alternatively, they will require the use of other targeted therapies to achieve tumor cell death.

Finally, we used our proteomic data identified protein complexes with potential therapeutic relevance in LGSOC. Using the CORUM database, we elucidated novel protein complexes differentially expressed between the MEKi response phenotypes. Previous studies have shown that some of these protein complexes regulate cell proliferation; including RFC in ovarian cancer [76] and MCM in other cancers [77,78]. The MCM complex forms the core of the DNA replicative helicase and, interestingly, is associated with cisplatin resistance [79]. Increased RFC expression has been shown to be associated with epithelial-mesenchymal transition and poor outcomes in both lung and breast cancer [80]. Other complexes we identified, such as the BRCC and the Condesin complexes, have been linked to DNA repair in cancer [81,82]. Although, MEKi represents a promising new therapy in LGSOC, MEKi treatment is only associated with a 6-8 months improvement in PFS. Increased expression of proteins within the complexes found in MEKi resistant cell lines provides a strong rationale to further explore these complexes as potential therapeutic targets.

LGSOC is a rare and often lethal cancer for which research model development has been challenging. As a result, our study is limited in the number of samples available for analysis. Data is not currently available to validate our findings in patients treated with MEKi. In this context, we have used multiomics analysis approaches as a disease driver and drug prediction biomarker discovery tool. Our study cohort has a higher frequency of p16 loss (chromosome 9p loss) than what is described in LGSOC tumors and thus models that preserve p16 expression will be needed for evaluation.

## Conclusion

Using a multiomics approach, we have characterized the largest worldwide collection of LGSOC cell lines to improve their use as research model systems. These cell lines accurately reflect the molecular make-up of LGSOC tumor samples characterized to date though LGSOC cell lines without 9p loss are not adequately represented in our sample. We have also described key molecular aberrations in our cell lines (including signaling pathway identification) that describe potential biomarkers of MEKi response and novel potential disease drivers. We found that proteomic data was more robust in differentiating MEKi drug sensitivity and identified several protein complexes that warrant further study in LGSOC. This approach has the potential to accelerate the discovery of clinically useful biomarkers and novel drug targets in MEKi-treated patient populations. Multiomics analytical approaches show great promise to improve our current understanding of cancer biology, drug therapy, and ultimately patient management.

## Additional files

Additional file 1: Supplementary material. (PDF 1.6 MB)

Additional file 2: Supplementary tables. (XLSX 1.3 MB)

## Abbreviations

AACR: American Association for Cancer Research;
AF: Allele frequencies;
BAM: Binary alignment map;
CNA: Copy number aberration;
CORUM: Comprehensive resource of mammalian protein complexes;
ER: Estrogen receptor;
FDR: False discovery rate;
GENIE: Genomics Evidence Neoplasia Information Exchange;
HGSOC: High-grade serous ovarian carcinoma;
HPLC: High performance liquid chromatography;
LGSOC: Low-grade serous ovarian carcinoma;
LOH: Loss of heterozygosity;
MCM: Minichromosome maintenance;
MEKi: MEK inhibitors;
MS: Mass spectrometry;
MSigDB: Molecular signature database;
OC: Ovarian Cancer;
PCR: Polymerase chain reaction;
PECA: Probe-level expression change averaging;
PFS: Progression free survival;
PKC-ɑ: protein kinase C alpha;
PR: Progesterone receptor;
RFC: Replication factor C-alpha;
RISC: RNA-induced silencing complex
SNV: Single nucleotide variation;
TMB: Tumor mutation burden;
TMT: Tandem mass tag;
UBC: University of British Columbia;
VAF: Variant allele frequency;
VGH: Vancouver General Hospital;
WES: Whole exome sequencing

## Acknowledgements

The authors thanks all members of the Carey’s and the Collin’s labs for helpful suggestions. The results published here are in part based upon data generated by the AACR Project GENIE https://www.synapse.org/#!Synapse:syn7222066.

## Funding

This study is funded by the Canada Foundation for Innovation (CFI-IF 33440), Terry Fox Research Institute (TFRI NF PPG Project #1062) (C.C.C), British Columbia Cancer Foundation (M.S.C) and the OvCaRe research program. The authors want to extend a special thanks to the MacKenzie and Lawler families, the Janet D. Cottrelle Foundation, and to all patients, families, and donors who supported this research. We would also like to thank the Society of Gynecologic Oncology of Canada (GOC) for their support in advancing research and knowledge translation in LGSOC, the Canadian Institutes of Health Research (CHIR) for their Planning and Disseminating grant support (NRF 152680), and the Women’s Health Research Institute (WHRI) for their Catalyst Grant support.

## Availability of data and materials

The whole exome and whole transcriptome sequencing data from this study is available in the European Genome-phenome Archive (EGA) under accession number XXXXXXX. The proteome data from mass spectrometry proteome data is available in the PRIDE Archive under accession number PXD019544.

## Authors’ contributions

RS, MLF, CCC, and MSC conceived the study and wrote the manuscript. RS, JH, SV, YL SA, BM, and RHB performed data analysis. ML, AD, HK, AH, SC, BC, SB, GED, and GBM performed the experiments generating the cell line models, specimen processing, quality control, sequencing, and mass spectrometry experiments. SLB, GBM, CCC, and MSC supervised the project, contributed scientific insights, and edited the manuscript. All authors read, contributed, and approved the final manuscript.

## Ethics approval and consent to participate

This study was approved by institutional human ethics review boards at BCCA (H14-02859), the University of British Columbia (UBC; R05-0119), and the University of Western Ontario (HSREB 12668E).

## Consent for publication

Not applicable.

## Competing interests

The authors declare that they have no competing interests.

## Notes

### Competing Interest Statement

The authors have declared no competing interest.

