## Supplementary materials for "Multiomics Characterization of Potential Therapeutic Vulnerabilities in Low-grade Serous Ovarian Carcinoma"

### Supplementary Material

### 1 Supplemental Methods

#### 1.1 Tumor samples and clinical information

Advanced or recurrent Low-grade serous ovarian Carcinoma (LGSOC) samples (tumor and ascites) were obtained from the OvCaRe gynecologic tumor bank (Vancouver General Hospital (VGH), British Columbia Cancer Agency, and the John and Mary Knight Translational Ovarian Cancer Research Unit (London Regional Cancer Program, London, Ontario, Canada). Tumor bank protocols, cell line derivation, and all research relating to this study was approved by institutional human ethics review boards at BCCA (H14-02859), the University of British Columbia (UBC; R05-0119), and the University of Western Ontario (HSREB 12668E). Clinical information was extracted retrospectively from patient records. Tumor histology was confirmed by a gynecological pathologist.

#### 1.2 Patient-derived LGSOC cell lines

LGSOC patient-derived cell lines were established through continuous in-vitro culture of patient material obtained through OvCaRe or the John and Mary Knight Translational Ovarian Cancer Research Unit (cell line iOvCa241) tumor banks (Fernández et al. 2016; Fernandez et al. 2019). Cultures were established and maintained in M199:MCDB105 (1:1) media (Cat. No. M5017 and M6395, Sigma-Aldrich, Oakville, Ontario, Canada) supplemented with 10% defined fetal bovine serum (dFBS; Cat. No. SH30070.03, Hyclone, GE Life Sciences, Logan, UT, USA) maintained at 37$^{o}$C and 5% CO2. No immortalization methods were used. Doubling time of these cells ranged from 30 to 80 h, with an average of 47 h, reflecting the clinical slow growth rate of LGSOC.

All cell lines were established from tumor tissues or ascites obtained from advanced/recurrent LGSOC patients, all in need of an effective systemic therapy. Eleven samples (**Supplementary Table 1**) were successfully established as a stable cell line, whereas three samples (CL-09, CL-11, and CL-12) were only grown as transient cell lines. In our previous studies (Fernández et al. 2016; Fernandez et al. 2019), which included a few of the cell lines above, we classified these cell lines according to their response phenotype to MEK inhibitor (MEKi) treatment. Cell lines CL-01, CL-02, CL-14, and CL-15 were classified as MEKi-sensitive lines after a single dose of MEKi treatment (100nM of trametinib, or 1 $\mu$M of selumetinib, or binimetinib, or refametinib) over a period of 4 days caused complete cell death. In contrast cell lines CL-03/CL-04, CL-05/CL-06, CL-07/CL-08 (three paired cell lines derived from consecutive tumors within three different patients) and CL-10 were classified as MEKi-resistant lines as they continued to proliferate under the same treatment conditions as above.

#### 1.3 Whole exome sequencing

DNA was extracted from LGSOC cell lines using an AllPrep DNA/RNA Mini Kit (Qiagen, Toronto, ON), and from matched patient buffy coat samples using a DNeasy Blood and Tissue Kit (Qiagen, Toronto, ON), according to the manufacturer's protocol. DNA concentration was quantified using a Qubit® dsDNA HS Assay (Thermo Scientific, Burlington, ON). About 0.5 $\mu$g of genomic DNA was fragmented by hydrodynamic shearing (Covaris, Massachusetts, USA) to generate 150-280 bp fragments. After end repair, fragments were ligated to Illumina barcoded adapters, cleanup, amplified by PCR to enrich for adapter ligated fragments, and controlled for quality. The library was then enriched by liquid phase hybridization using Agilent SureSelect XT Human All Exon v6 (Agilent Technologies, CA, USA) following the manufacturer's recommendations. The captured library was amplified by PCR using indexing primers, cleanup and quality assessment was done with the TapeStation 4200 (Agilent). Libraries are submitted to PE100 sequencing on Illumina HiSeq4000.

##### 1.4 Somatic variant calling

Sequence alignment and mutation calling were performed in Partek Flow environment (© Partek Inc). Sequence reads were aligned to the GRCh38/hg38 human genome build using bwa 0.7.2 (Li and Durbin 2009) Variants were called using Strelka 1.0.15 (Kim et al. 2018) for all cell lines except for CL-02 (lacking buffy coat sample). CL-02 variant calling was performed using LoFreq 2.1.3.a (Wilm et al. 2012). The called variants were annotated using the Annovar software (Wang et al. 2010). Annotated calls were then filtered to show only protein-changing SNVs that were present in cell line DNA at allele frequencies (AF) greater than 0.1 and coverage higher than 16x. For CL-02, all calls not present in dbSNP (version 138) were retained, while of the calls that were present in dbSNP, calls with (average heterozygosity + aveHet standarderror) less than 0.1 were retained. These were additionally filtered using the same criteria as for the Strelka calls.

##### 1.5 Copy number aberration calls

Data analysis was performed using Nexus Copy Number Discovery Edition Version 9.0 (BioDiscovery, Inc., El Segundo, CA). Samples were processed using the Nexus NGS functionality with the FASST2 segmentation. The log ratio thresholds for single copy gain and single copy loss were set at +0.18 and -0.18, respectively. The log ratio thresholds for gain of 2 or more copies and for a homozygous loss were set at +0.6 and -1.0, respectively. Tumor sample BAM files were processed with corresponding normal tissue BAM files. Probes were normalized to the median.

##### 1.6 Whole transcriptome sequencing (RNA-seq)

Total RNA from was isolated in accordance with the mirVana miRNA Isolation kit (ThermoFisher). Strand specific RNA sequencing was performed on quality controlled high RIN value (>7) RNA samples (Bioanalyzer Agilent Technologies). In brief, 200 ng of total RNA was first treated to remove the ribosomal RNA (rRNA) and then purified using the Agencourt RNA Clean XP Kit (Beckman Coulter) prior to analysis with the Agilent RNA 6000 Pico Chip to confirm rRNA removal. Next, the rRNA-depleted RNA was fragmented and converted to cDNA. Subsequent steps include end repair, addition of an 'A' overhang at the 3' end, ligation of the indexing-specific adaptor, followed by purification with Agencourt Ampure XP beads. The strand specific RNA library prepared using TruSeq stranded mRNA kit (Illumina). Libraries were quality checked and sized with a TapeStation 4200 (Agilent Technologies), then quantified using the KAPA Library Quantification Kit for Illumina platforms (Kapa Biosystems). Libraries were submitted to PE75 sequencing on Illumina NextSeq500. About 20-50 million sequencing reads per library was analyzed for each sample.

##### 1.7 Transcriptome (RNA-seq) quantification

RNA-seq sequence alignment and transcript/gene quantification were performed in Partek Flow environment (© Partek Inc). The RNA-seq reads were first trimmed to remove low quality bases from the 3' end. The trimmed reads were then mapped to the GRCh38/hg38 human reference genome and transcriptome (RefSeq release 83) using splice-aware aligner STAR (2.5.3a) (Dobin et al. 2013). Based on the reads that can only be mapped to a single genomic location, the transcript/gene expression quantification was performed using HTSeq-count (Anders et al. 2015). Cross-sample normalization of expression values were done by DESeq (Anders and Huber 2010).

##### 1.8 Proteomics analysis using mass spectrometry

Seven LGSOC cell lines were selected for global proteome analysis (**Supplementary Table 1**) in triplicates using 3 separately cultured samples for each. Cultures of 5 different HGSOC cell lines (OVCAR8, OVSAYO, CAOV3, OVCAR4, and VOA2900) were also used as a control. VOA2900 is a HGSOC cell line developed by the ovarian cancer research program at VGH. Cells were collected by trypsinization and washed with 1X dPBS. A minimum of $1X10^6$ cells per cell line were used in these analyses. Cells pellets were lysed in 200 $\mu$L of a buffer containing guanidine hydrochloride (4M), HEPES (pH 8.5, 50 mM), 2-chloroacetamide (40 mM), tris(2-carboxyethyl) phosphine (10 mM), ethylenediaminetetraacetic acid (5 mM), 1x phosphatase inhibitor, and 1x protease inhibitor. Bead beating of the cells was performed in Lysing Matrix D tubes on a FastPrep24 (6 M/s for 45 s, 60 s rest, 6 M/s for 45 s). Cell lysate was then heated at 95$^o$C for 15 min with 1200 rpm mixing. 50 $\mu$g of each sample was diluted 10x in 50 mM HEPES pH 8.5 (concentration determined using by BCA assay) and digested using a 50:1 ratio of protein:trypsin/Lys-C mixture at 37$^o$C overnight with shaking at 1200 rpm. A small aliquot of each LGSOC sample was mixed and analyzed as LGSOC-mixed control. The five HGSOC samples were mixed and used as a HGSOC-mixed control sample. An aliquot of all the LGSOC and HGSOC samples were mixed and used as an all sample pooled internal standard. Tandem mass tag (TMT10) labeling was performed by adding 100 $\mu$g TMT label in acetonitrile and incubating at room temperature for thirty minutes twice. Excess label was quenched with 10 $\mu$L of 1 M glycine. Individual TMT channels were combined and the volume of the combined TMT labeled sample was reduced to 10-20% of the original volume. Three 10-plex batches were prepared, where each batch contained one replicate of each of the LGSOC samples, the LGSOC-mixed control, the HGSOC mixed control, and the all sample pooled internal standard. The combined samples were fractionated by high pH reverse-phase high performance liquid chromatography (HPLC) in a gradient of acetonitrile and 20 mM ammonium bicarbonate in water into 48 individual fractions. These 48 fractions were concatenated into 12 fractions by combining every twelfth fraction, each of which was analyzed by low pH nanoLC mass spectrometry (MS) using a MS/MS/MS (MS3) method on a Thermo Scientific Easy nLC coupled to a ThermoFisher Orbitrap Fusion MS. The Uniprot Human Proteome was used as a reference proteome (UniProt Consortium 2018). Only peptides that were identified in all three batches were retained. ComBat, a function from sva R-package (Leek et al. 2012), was used to adjust for batch effects using an empirical Bayes adjustment. Detection of differentially expressed proteins from the collected peptide data was determined by the Probe-level Expression Change Averaging (PECA) in R (Teo et al. 2014).

##### 1.9 Mutational signature analysis

We used deconstructSigs (Rosenthal et al. 2016), a multiple regression approach to statistically quantify the contribution of mutational signature for each tumor. The 30 mutational signatures were obtained from the COSMIC mutational signature database (Forbes et al. 2017). In brief, deconstructSigs attempts to recreate the mutational pattern using the trinucleotide mutation context from the input sample that closely resembles each of the 30 mutational signatures from COSMIC mutational signature database (v2 - March 2015) (Forbes et al. 2017). In this process, each mutational signature is assigned a weight normalize between 0 to 1 indicating its contribution. Only those mutational signatures with a weight more than 0.06 were considered for analysis. We further combined (or grouped) these 30 mutational signatures based on the similarity of their mutational features or functional associations.

- Mutation signatures 3, 6, and 15 are related to DNA mismatch repair, so they were grouped together.
- Mutation signatures 5 and 12 exhibits strong transcriptional strand bias for T>C mutations, so they were grouped together.
- Mutational signatures 8, 24 and 29 were grouped together because they exhibit transcriptional strand bias for C>A mutations indicating guanine damage that is most likely repaired by transcription-coupled nucleotide excision repair.

##### 1.10 Prioritization of driver genes using HIT'nDRIVE

Using our recently developed computational algorithm HIT'nDRIVE (Shrestha et al. 2017), we identified driver genes in these LGSOC cell lines. Briefly, HIT'nDRIVE measures the potential impact of genomic aberrations on changes in the global expression of other genes/proteins which are in close proximity in a gene/protein-interaction network. It then prioritizes those aberrations with the highest impact as cancer driver genes. Both non-silent somatic mutation calls and CNA gain or loss were independently collapsed in gene-patient alteration matrix with binary labels. mRNA and protein expression values were used to derive expression-outlier gene-patient outlier matrix using Generalized ESD Many-Outlier Deviate (GESD) test (Rosner 1983). STRING ver10 (Szklarczyk et al. 2014) protein-interaction network was used to compute pairwise influence value between the nodes in the interaction network. We integrated these genome and transcriptome as well as genome and proteome data using HIT'nDRIVE algorithm. First we integrated genome and transcriptome data. Here we ran HIT'nDRIVE separately using SNV-mRNA expression data and CNA-mRNA expression data. For this we used the parameters: $\alpha = 0.9$, $\beta = 0.7$, and $\gamma = 0.7$. Next, we integrated genome and proteome data. Here we ran HIT'nDRIVE separately using SNV-protein expression data and CNA-protein expression data. For this we used the parameters: $\alpha = 0.9$, $\beta = 0.7$, and $\gamma = 0.8$. We used IBM-CPLEX as the Integer Linear Programming solver. The results were later combined to downstream analysis.

##### 1.11 Ordination analysis

First, for mRNA expression data, we selected the expression profile of protein coding genes only. We performed median absolute deviation (MAD) analysis independently on mRNA expression and protein expression data. We then selected those genes/proteins whose MAD value exceeded the 75th percentile of the MAD values of all genes/proteins for further analysis. We then performed ordination using the principal components analysis (PCA). For this we used *prcomp()* function of R *Stats* Package.

##### 1.12 Differential expression analysis

Differential expression analysis of MEKi-response phenotypes was performed by applying linear empirical Bayes model using "sva" R-package (Leek et al. 2012). Gene (mRNA) and protein expression data of the MEKi-response phenotypes were used for this purpose. We used the following threshold values to select genes and proteins for MEKi response analysis and downstream pathway analysis: for mRNA expression data, $pvalue \leq 0.05$ and $|foldchange| > 1.03$, and for protein expression data, $pvalue \leq 0.05$ and $|foldchange| > 1.03$. Furthermore, to identify differentially expressed protein complex, we used Wilcoxon rank-sum test on the average protein expression profiles of protein complex members. Protein complexes with $pvalue \leq 0.05$ and $|foldchange| > 1$ were selected differential protein complex analysis.

##### 1.13 Pathway enrichment analysis

The differentially expressed set of genes/proteins were tested for enrichment against gene sets of KEGG pathways present in the Molecular Signature Database (MSigDB) v6.0 (Subramanian et al. 2005). A hypergeometric test based over-representation analysis was used for this purpose (https://github.com/raunakms/GSEAFisher). A cut-off threshold of FDR$\leq$ 0.01 was used to obtain the significantly enriched pathways. Only pathways that are enriched with at least four differentially expressed genes were considered for further analysis. To calculate the pathway activity score, the expression dataset was transformed into standard normal distribution using "inverse normal transformatio" method. This step is necessary for a fair comparison between the expression-values of different genes. For each sample, the pathway activity score is the mean expression level of the differentially expressed genes linked to the enriched pathway. The KEGG signaling pathways and associated expression profile were visualized using pathview R-package (Luo and Brouwer 2013).

##### 1.14 External datasets

We utilized DNA sequencing datasets of publicly available patient LGSOC cohort from the American Association for Cancer Research (AACR) project Genomics Evidence Neoplasia Information Exchange (GENIE) AACR Project GENIE Consortium (2017). The dataset consisted of a total of 122 LGSOC tumors (both primary and metastatic tumors). We used somatic mutation and copy number aberration profiles from the dataset. AACR GENIE Project Data: Version 5.0 was downloaded from (SynapseID: syn7222066) https://www.synapse.org/Synapse:syn7222066.

##### 1.15 Protein complex co-expression score

For a given protein complex, its co-expression score is computed as the average Pearson correlation coefficient of all pairwise protein-protein interactions (as represented in STRING ver10 (Szklarczyk et al. 2014) database) in the complex. The co-expression scores were separately calculated using mRNA expression data ($R_{mRNA}$) and protein expression data ($R_{protein}$). Protein complexes with ($|R_{mRNA} - R_{protein}| \leq 0.05$) are defined as correlated protein complexes. For this analysis, we considered the CORUM (Ruepp et al. 2009) protein complexes consisting of at least 4 proteins.

### Supplemental Figures

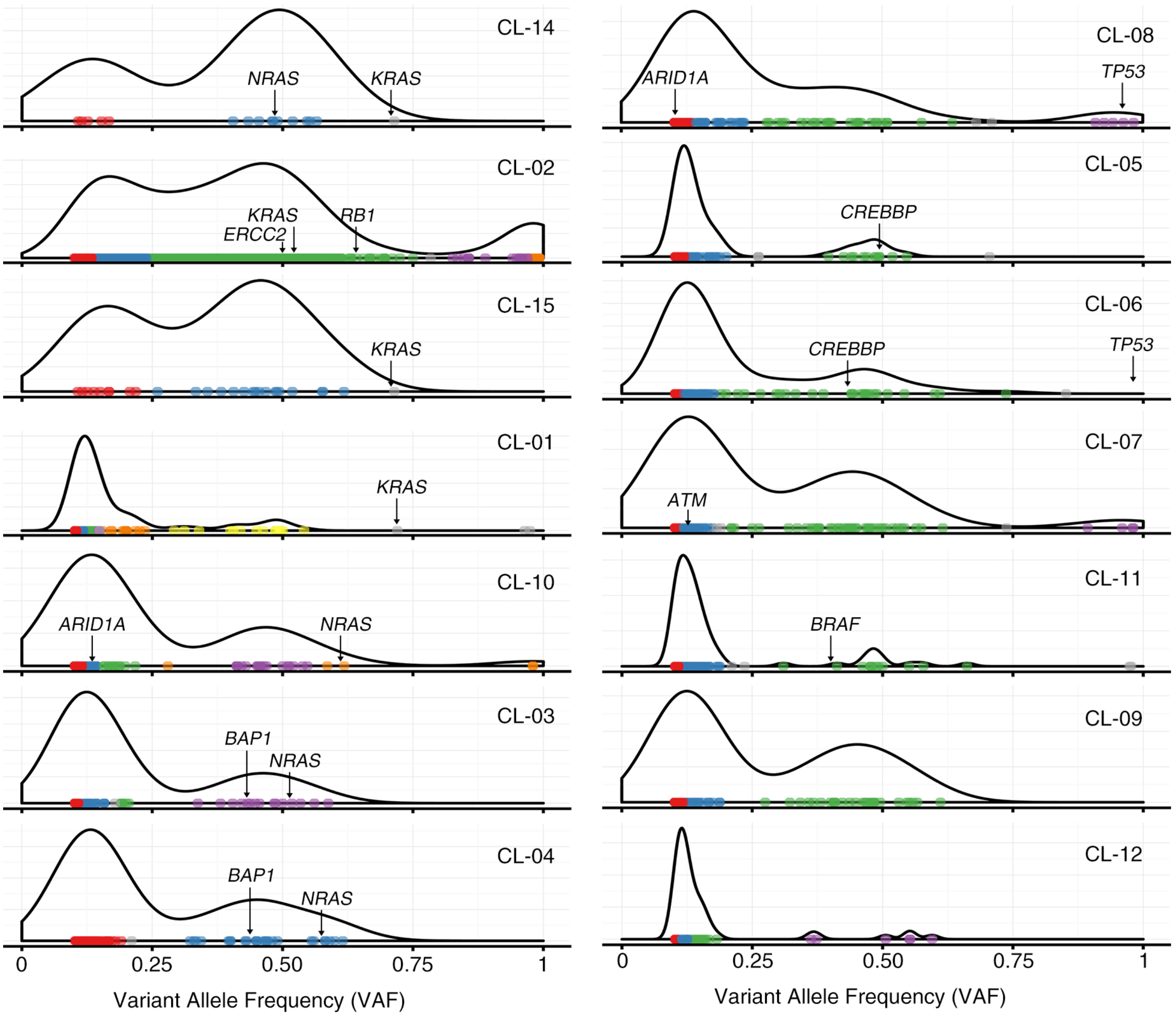

**Supplementary Figure 1. Distribution of variant allele frequency (VAF) of mutations in LGSOC.** Based on VAF, the somatic mutations identified in LGSOC were clustered into different groups using the R-package Maftools. The horizontal axis represents the VAF of the mutations and the vertical axis represents its density.

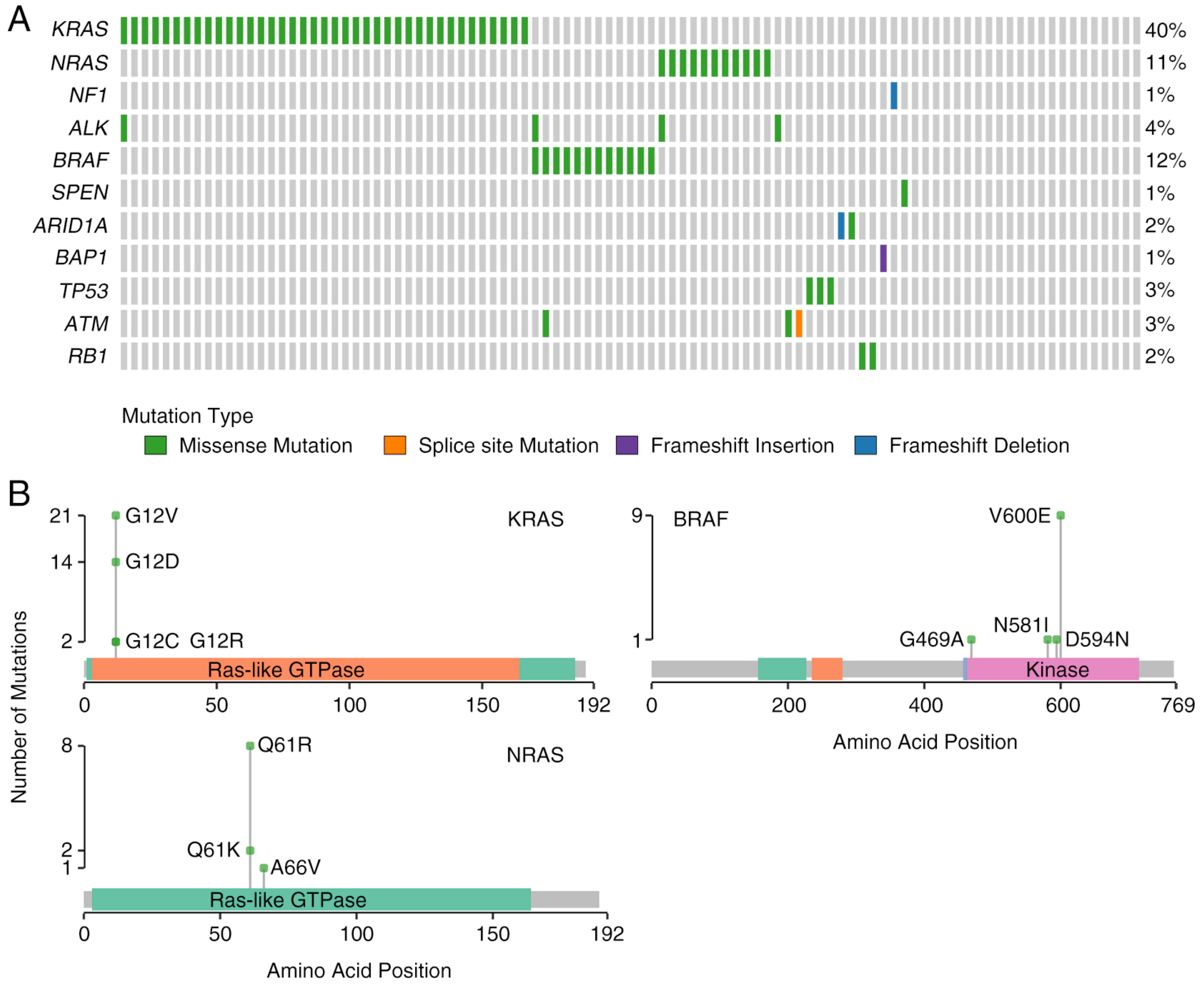

**Supplementary Figure 2. Somatic Mutation Landscape of LGSOC in AACR Project GENIE Cohort.** (A) Oncoplot showing the status of mutated genes in major cancer pathways - MAPK pathway, Notch pathway, chromatin remodeling, and DNA repair pathway. Genes that overlap with those shown in Figure 1A is represented here. The numbers on the right of the plot shows the recurrence frequency (in percentage) of the corresponding mutation in the cohort. (B) Plots showing mutation distribution and the protein domains for the corresponding mutated protein.

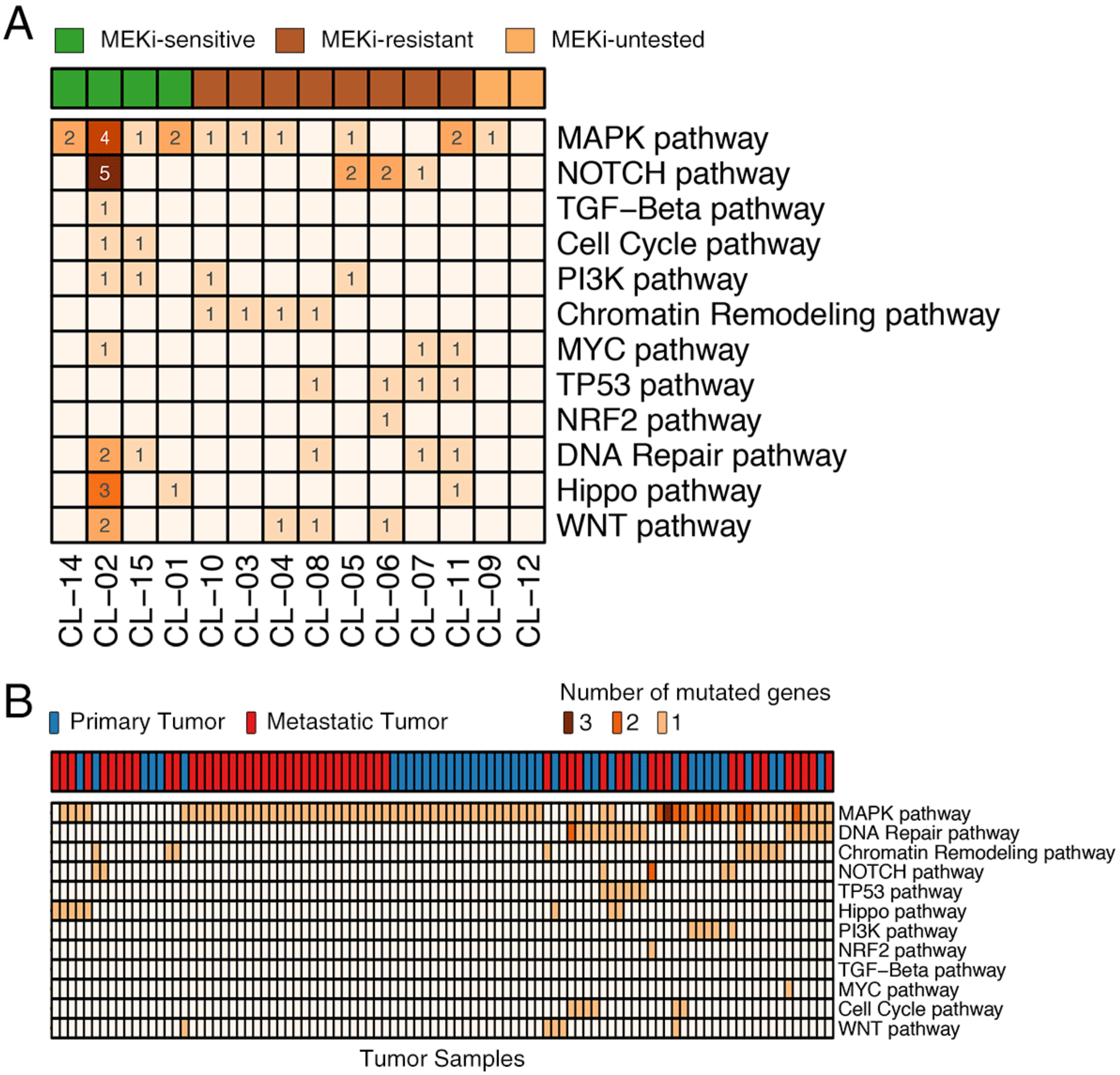

**Supplementary Figure 3. Mutated Oncogenic Pathways in LGSOC.** (A) Heatmap showing the number of genes in the respective oncogenic pathways that are mutated per LGSOC cell line. The number of genes mutated in each pathway is indicated. (B) Mutated oncogenic pathways in LGSOC tumors from AACR Project GENIE. Heatmap showing the number of genes in the respective oncogenic pathways that are mutated per LGSOC tumor.

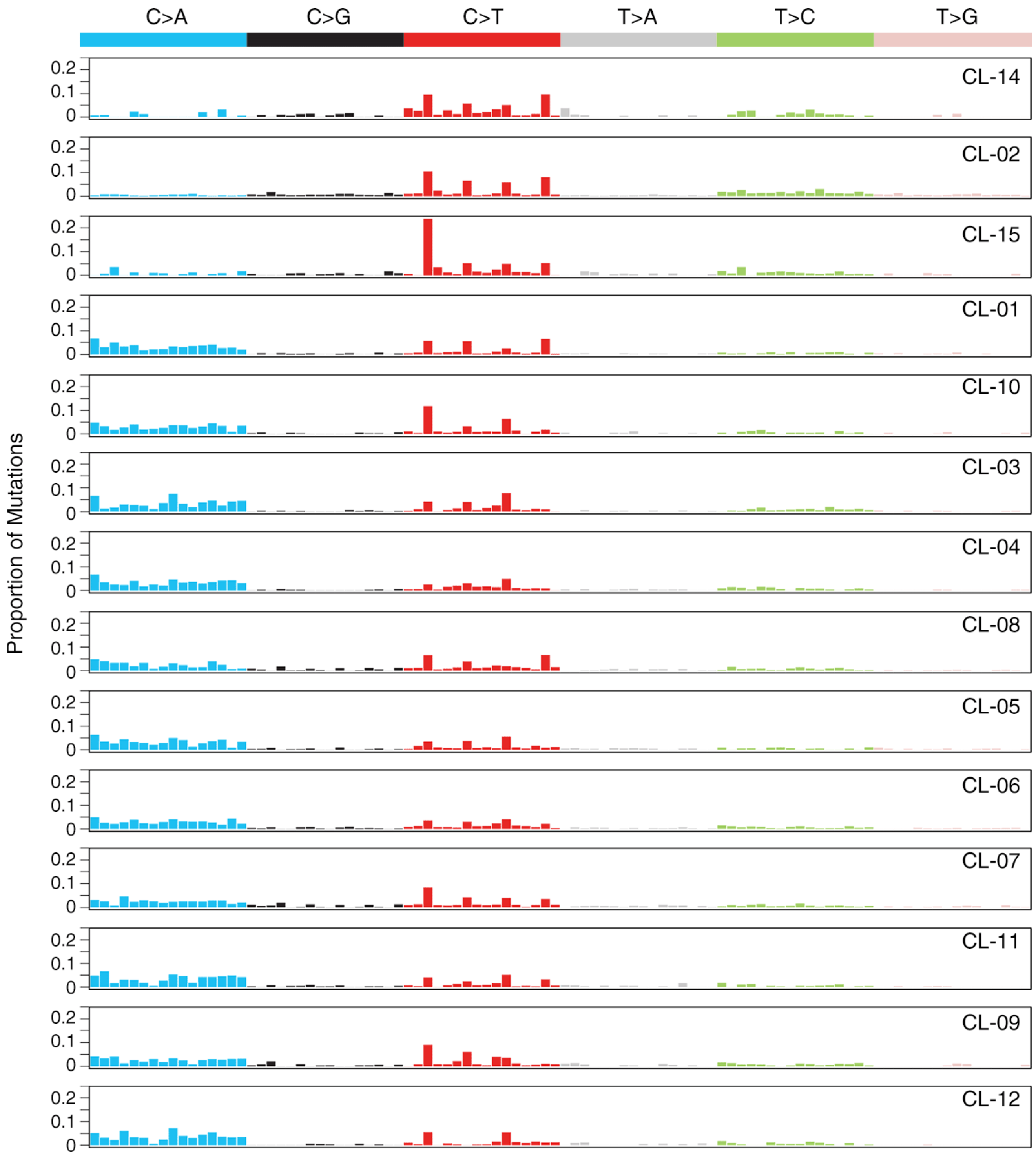

**Supplementary Figure 4. Nucleotide Substitution Mutation Patterns in LGSOC.** The horizontal axis represents each of the 96 different possible combinations of the tri-nucleotide substitution mutation combinations colored by their respective substitution patterns. The vertical axis represents the proportion of mutations per sample.

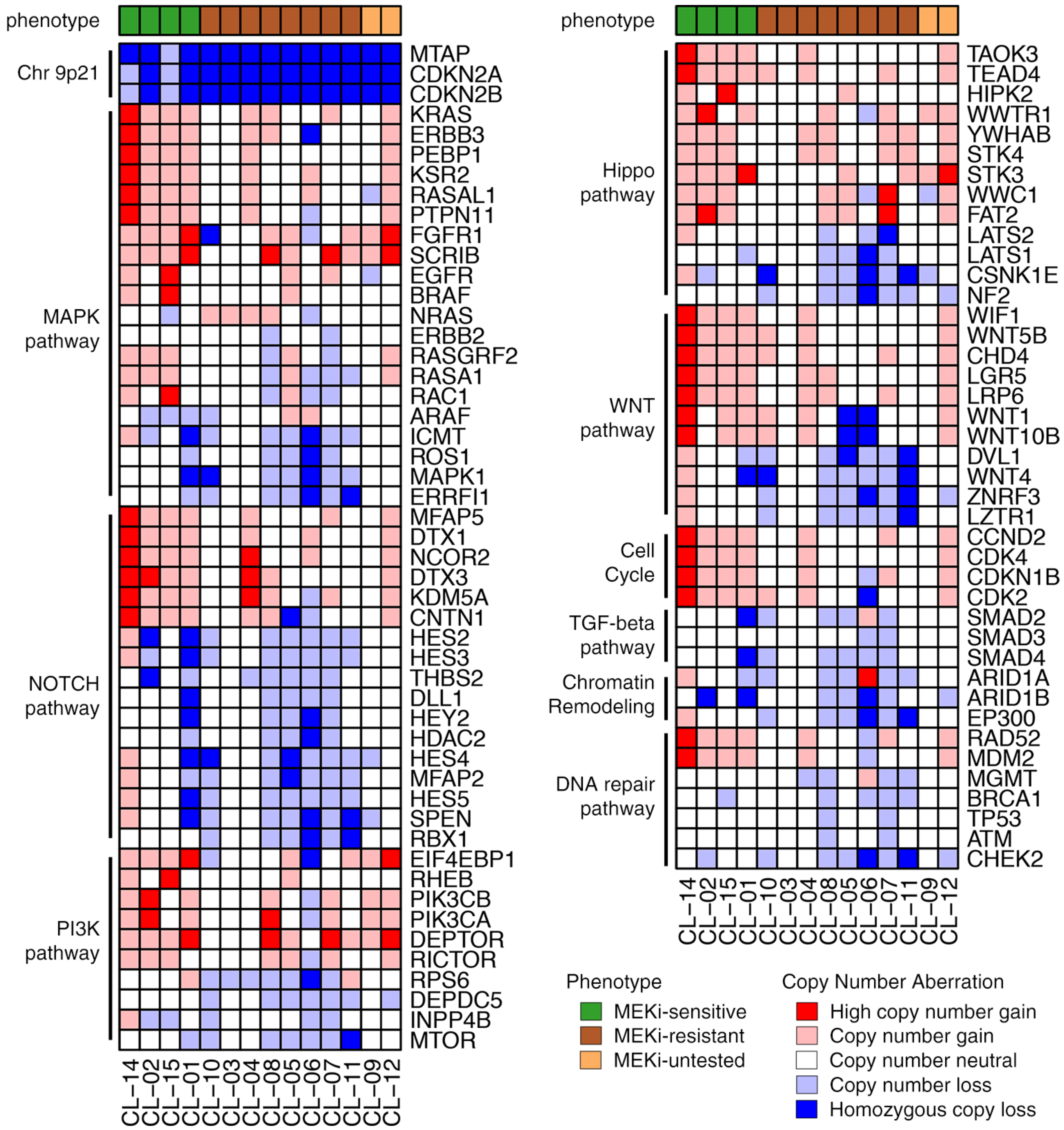

**Supplementary Figure 5. Copy number aberration status of LGSOC cell lines in oncogenic pathways.** Heatmap showing the copy number status of genes grouped by different oncogenic pathways. In addition, genes in chromosome 9p21 are also shown.

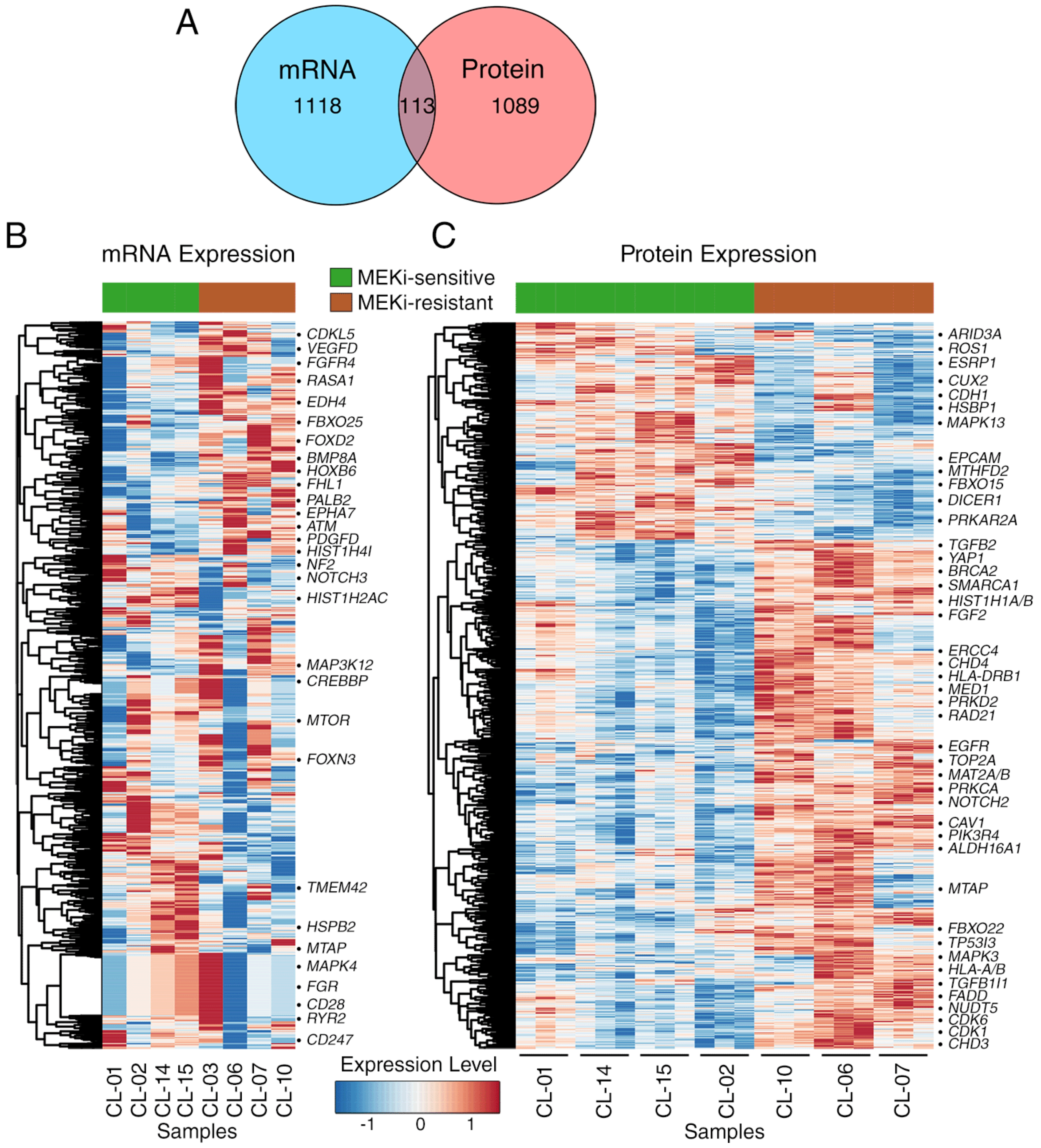

**Supplementary Figure 6. Significantly differentially expressed genes/proteins between the MEKi drug response phenotypes.** (A) Venn diagram of overlap between the differentially expressed genes (from mRNA expression profiles) and proteins (from protein expression profiles) between the MEKi drug response phenotypes. (B-C) Heatmap of mRNA and protein expression profiles of the significantly differentially expressed genes/proteins between the MEKi- drug response phenotypes. The expression profile of each gene were mean normalized for visualization. Key differentially expressed genes have been highlighted.

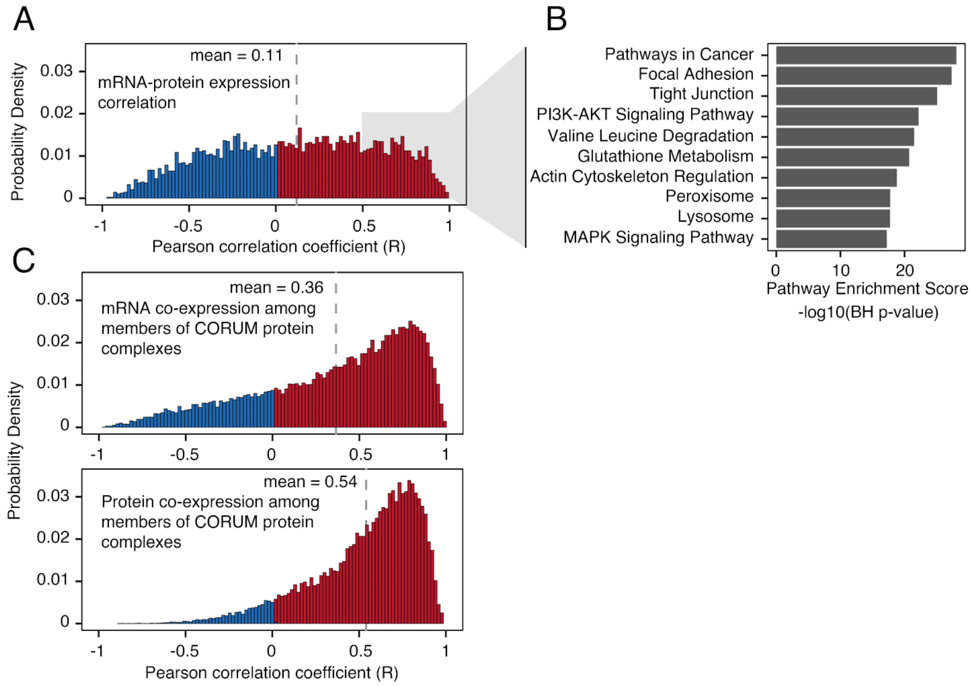

**Supplementary Figure 7. mRNA and protein correlation.** (A) Histogram showing the distribution of Pearson correlation coefficient (R) of mRNA and protein expression correlation in LGSOC cell lines. Around 25% (1271 out of 4982) of proteins were highly correlated ($R \geq 0.5$) with their corresponding transcript abundance. (B) KEGG pathway enrichment of genes with high mRNA and protein expression correlation ($R \geq 0.5$). These highly correlated proteins were involved in different oncogenic signaling pathways including Focal adhesion, PI3-AKT pathway, and MAPK pathway. (C) Histogram showing the distribution of Pearson correlation coefficient (R) of mRNA co-expression and protein co-expression among the members of CORUM protein complexes. We compared the transcript abundances (co-expression) of a number of protein complex members. About 48% (24636 of 51275 gene pairs in 1701 protein complexes) of all gene pairs measured were highly correlated ($R \geq 0.5$). We also compared the protein abundances of these protein complex members. Interestingly, the protein abundance of protein complex members were highly co-expressed as compared to their respective transcripts. About 63% (22546 of 35411 protein pairs in 1008 protein complexes) of all protein pairs measured were highly correlated ($R \geq 0.5$).

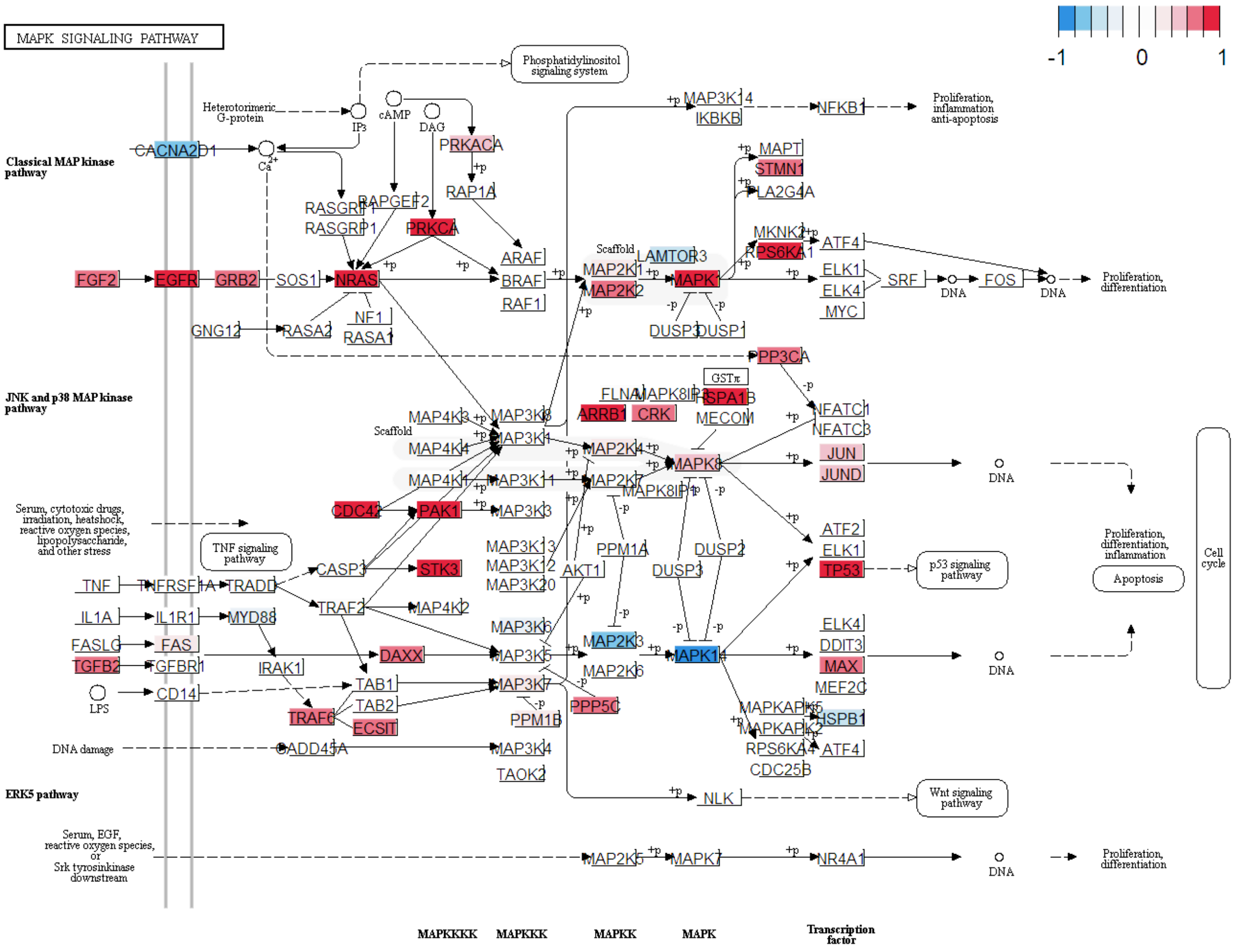

**Supplementary Figure 8. MAPK Signaling Pathway in MEKi-resistant LGSOC cell lines.** Schematic overview of MAPK Signaling Pathway obtained from the KEGG pathway database. Individual proteins identified in the mass spectroscopy experiment were mapped into MAPK Signaling Pathway and their average protein expression profile in MEKi-resistant cell lines were visualized using *pathview* R-package.

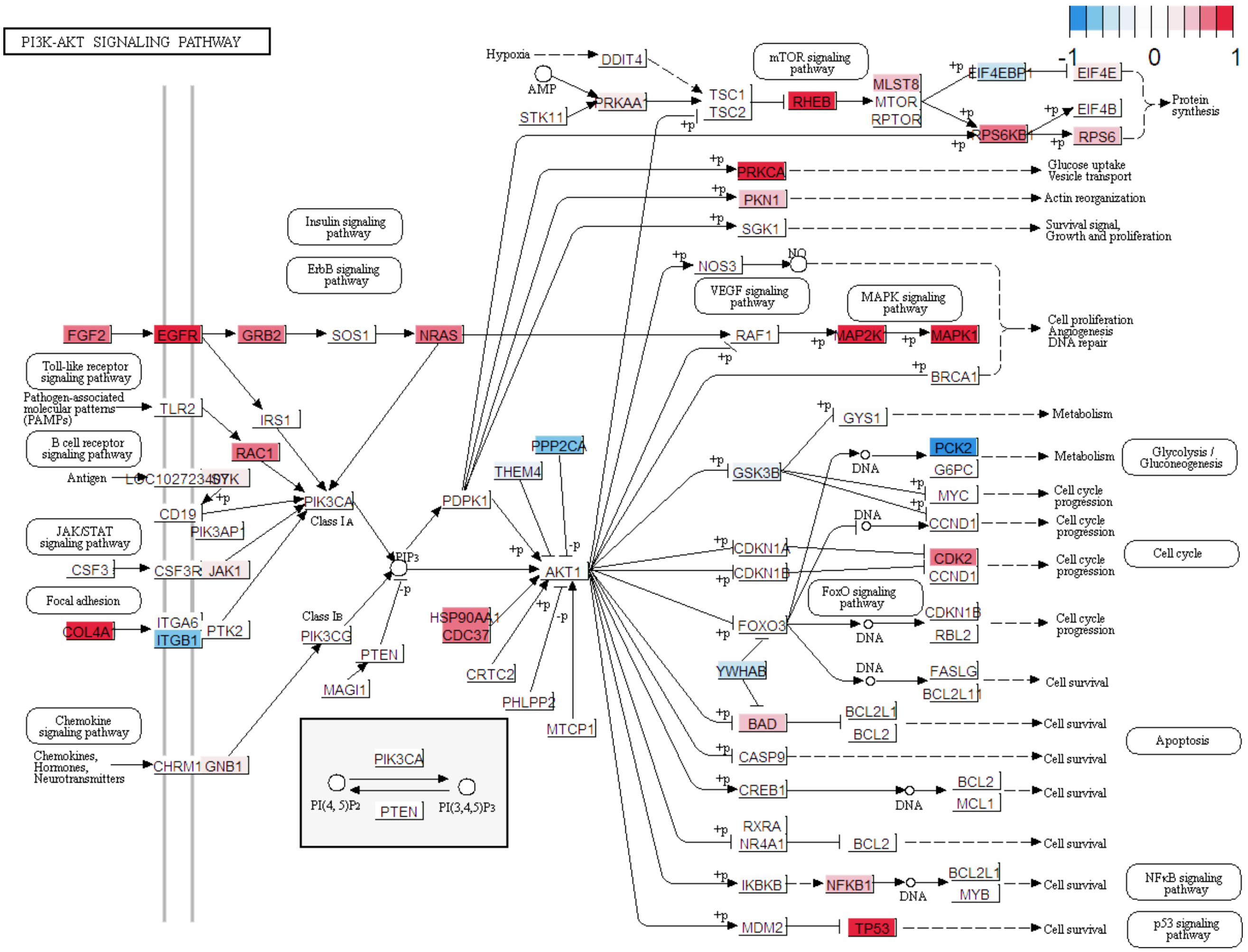

**Supplementary Figure 9. PI3K-AKT Signaling Pathway in MEKi-resistant LGSOC cell lines.** Schematic overview of PI3K-AKT Signaling Pathway obtained from the KEGG pathway database. Individual proteins identified in the mass spectroscopy experiment were mapped into PI3K-AKT Signaling Pathway and their average protein expression profile in MEKi-resistant cell lines were visualized using *pathview* R-package.
